# A new universal chimeric-antigen receptor (CAR)- fragment antibody binder (FAB) split system for cancer immunotherapy

**DOI:** 10.1101/2024.12.05.627079

**Authors:** Ainhoa Arina, Edwin Arauz, Karolina Warzecha, Elham Masoumi, Annika Saaf, Łukasz Widło, Tomasz Slezak, Aleksandra Zieminska, Karolina Dudek, Zachary P. Schaefer, Maria Lecka, Svitlana Usatyuk, Ralph R. Weichselbaum, Anthony A. Kossiakoff

## Abstract

Chimeric antigen receptor T (CAR-T) cell therapy has shown extraordinary results in treating hematological cancer. However, many patients relapse because of heterogeneous antigen expression and outgrowth of antigen lost variants. Other problems include on-target-off-tumor toxicity and the requirement for manufacturing of complex cellular products. Universal and modular CAR constructs offer significantly improved flexibility, safety and cost-effectiveness over conventional CAR constructs. Here we present a new chimeric-antigen receptor (CAR)-fragment antibody binder (Fab) platform based on an engineered protein G variant (GA1) and Fab scaffolds that present exquisite specificity and selectivity on antibody capture. The expression of GA1CAR on human CD8^+^T cells leads to antigen recognition and T cell effector function that can be modulated according to the affinity of the CAR for the Fab scaffold and of the Fab for the target. GA1CAR-T cells can recognize multiple Fab-antigen pairs on breast and ovarian cancer cell lines. Adoptive transfer of GA1CAR-T cells/Fabs in breast cancer xenograft models leads to effective tumor control. Rapid re-direction of the CAR-T cells to a new target can be achieved by using different Fabs. GA1CAR expression confers favorable phenotypic properties to T cells including a higher effector function upon exposure to antigen as compared to conventional scFv CAR-T cells. This highly versatile “plug and play” CAR-T platform has potential for application in personalized therapy, preventing antigen loss variant escape, decreasing toxicity and increasing access.

## Introduction

Chimeric antigen receptor T (CAR-T) cell therapy has demonstrated impressive success in the treatment of hematological malignancies, such as acute lymphoblastic leukemia and non-Hodgkin lymphoma, by genetically engineering T lymphocytes to target specific cancer antigens. Despite these advances, CAR-T therapy faces several significant challenges, including low tumor penetration, severe toxicities like tumor lysis syndrome, cytokine release syndrome, and CAR-T related encephalopathy syndrome [1–3]. Many patients experience relapse due to antigen heterogeneity or loss [4–10], and the therapy’s application to solid tumors has been limited by factors such as low infiltration into the tumor microenvironment and high antigenic variability [6], [11–16]. Many adverse side effects stem from potent off target effects that induce killing of normal tissue that expresses a threshold amount of the target protein [17–18]. Current CAR-T treatments are highly personalized and complex, leading to high costs and limited accessibility. To address these issues, innovative strategies are needed to enhance safety, efficacy, and accessibility.

Effective CAR-T therapy depends on manifold antigen engagement properties like antigen density, inter-membrane distance, epitope location and orientation, and targeting moiety (scFv), as well as its spacer length [19–20]. Considering the number of interrelated parts, it is clear that no standard template can be followed to construct the optimal CAR- T candidate. Factoring the timing considerations and cost of production, it is virtually impossible to undertake patient specific optimization of traditional CAR-T constructs. With these considerations in mind, numerous efforts have been made to develop “universal” CAR-T constructs based on modular, split, and switchable systems that allow precise control of the cell therapy’s timing, strength and specificity [21]. These systems cover a range of quite different approaches [21–26]. However, these approaches still often involve complex designs with structural constraints.

To address this issue, described here is the development and testing of GA1CAR, a novel modular antigen receptor system that utilizes pairing an engineered protein G variant (GA1) with a set of antibody fragment Fabs with different modifications in their scaffolds. In this system, the Fab module has dual functions: antigen recognition through the CDR loops of its variable domain and attachment to GA1 of the CAR through its constant domain. Importantly, the affinities of either contact interface can be adjusted to affect the binding dynamics of GA1CAR without altering the targeting of the tumor antigen or GA1. This design allows for targeting multiple antigens by simply swapping out the particular Fab scaffold, offering a “plug-and-play” approach that enhances versatility in targeting different tumor antigens [27–28]. The GA1CAR system allows for affinity-dependent modulation of effector functions. By using modified Fab scaffolds having varying affinities to GA1, the GA1CAR can fine-tune therapeutic responses such as cytokine and IFNγ release, providing a mechanism to adjust the intensity of the immune response. Further, the interchangeable Fab scaffolds provide an on-off switching capability and provide for the ability to assess the patient’s tolerance to the GA1CAR in a systematic way, exploiting potency differences that couple with the different Fab scaffold affinities to GA1CAR.

Using xenograft mouse models, we determined the in vivo efficacy of GA1CAR-T cells and the ability to redirect GA1CAR-T cells from one target to another by treating the animals with Fabs targeting different tumor-specific antigens. Comparison of the phenotype of GA1 vs conventional scFv CAR-T cells showed a higher activation and effector function upon contact with cancer cells for the GA1CAR-T, together with lower tonic signaling gene expression at baseline. Additionally, we show that the unloaded GA1CAR-T can remain dormant in the animal for an extended period and then, in the case of tumor rebound, be reloaded by reinfusing the target appropriate Fab. Taken together, this novel modular CAR system, the GA1CAR, is endowed with higher versatility, flexibility and safety advantages compared to standard scFv CARs, and lower complexity than other modular CARs. It has demonstrated specificity for various antigens overexpressed in breast and ovarian cancers and shows promise in preclinical models for personalized therapy applications.

## Results

### Design and Generation of GA1CAR

The GA1CAR system is a modular chimeric antigen receptor that utilizes a specially engineered protein G variant, known as GA1, along with engineered Fab scaffolds developed through phage-display technologies **(Fig. 1A)**. Protein G naturally binds to immunoglobulins, but the GA1 variant has been engineered to prevent binding to naturally occurring antibodies and to eliminate potential immunogenicity [29]. The Fab scaffolds are based on the Herceptin Fab framework (4D5), which is widely used in clinical applications due to its stability and low immunogenicity [29]. This Fab framework has been further modified to create LRT, SQRT, or wild-type (wt) Kappa scaffolds by introducing specific mutations into the Fab’s constant light chain [27–28]. The interaction affinities between these Fab scaffolds and GA1 were measured using Surface Plasmon Resonance (SPR), revealing dissociation constants (Kd) of 100 pM for LRT, 10 nM for SQRT, and 50 nM for 4D5. Fabs with wt human Kappa light chains do not bind to GA1 and serve as negative controls [27].

**Fig.1.**
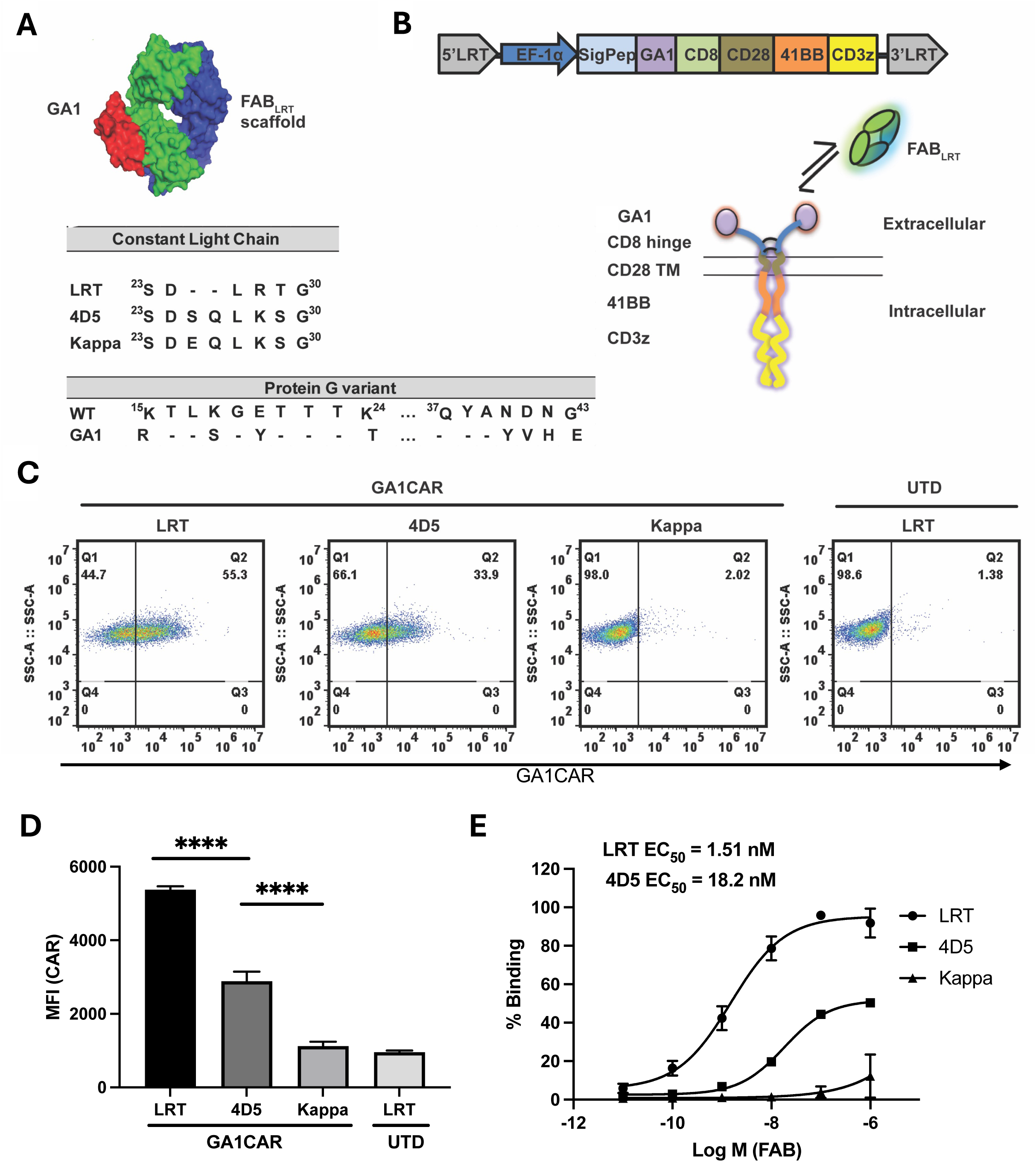
Design of the GA1CAR. A) Cartoon showing binding of protein GA1 (red) and Fab^LRT^ scaffold (green for Lc, blue for Hc); as well as engineered mutations present on different Fab scaffolds and protein GA1. B) Schematic of lentiviral vector for GA1CAR containing protein GA1, CD8 hinge, CD28 TM, 41BB, and CD3ζ domains; and cartoon of GA1CAR on the cell surface and its Fab capturing mechanism. C) Representative plots showing the surface expression of GA1CAR in Jurkat cells after lentiviral transduction, determined by flow cytometry after incubation with an irrelevant Fab^LRT^ followed by a secondary anti-human Fab antibody conjugated to Alexa Fluor® 647. D) Quantification by flow cytometry of different Fab scaffolds binding to GA1CAR expressed on Jurkat cells; detection was done by staining with an anti-human Fab antibody conjugated to Alexa Fluor® 647. Fabs were used at 200 nM. E) EC50 determination of different Fab scaffolds binding to GA1CAR-J, The data are presented as the mean ± SD of four independent experiments (n = 4). Statistical significance was determined by Tukey’s multiple comparisons test after one-way ANOVA (*P < 0.05; **P < 0.01; ***P < 0.001; ****P < 0.0001), P < 0.05 considered significant.

The schematic of the GA1CAR lentivirus construct and its modular components to form a second generation CAR assembly are illustrated in **Fig. 1B**. The GA1 module was fused upstream of a CAR-T construct containing a CD8 hinge, CD28 transmembrane domain, 41BB co-stimulatory domain, and CD3ζ domain (**Extended Data Fig 1**). The production of a functional GA1CAR was confirmed through lentiviral transduction in Jurkat cells (GA1CAT-J), using Fabs that bind the GA1 module, followed by staining with a fluorescent anti-human Fab’2 antibody **(Fig. 1C**). The LRT scaffold showed superior binding to GA1 compared to the others **(Fig. 1D-E**), consistent with the observed dissociation constants. A single molecule pulldown assay was used to verify the dimerization of the CD8 hinge in the GA1CAR structure (**Extended Data Fig 2**). Using three distinct pulldown strategies - anti-SNAP antibody (**Extended Data Fig 2A**), biotinylated peptide antigen (**Extended Data Fig 2B**), and anti-YFP pulldown (**Extended Data Fig 2C**) - we counted immobilized molecules and their photobleaching steps. The resulting distribution, predominantly showing one and two photobleaching steps, supports a dimeric stoichiometry for the CAR [30–31].Together, this data confirms successful generation and expression on human T-cell surfaces.

### Functional Characterization of GA1CAR

To assess how affinity affects functional response amplitude towards a surface expressed antigen, we use HEK cells expressing maltose binding protein (MBP) and a conformationally sensitive anti-MBP Fab engineered with an LRT scaffold (Fab^LRT^). This Fab was generated by phage display to bind to the surface-expressed MBP in a manner that depended on the concentration of added maltose **(Extended Data Fig 3)** [32] .Thus, the GA1CAR/anti-MBP Fab system allowed for the evaluation of effector cell activation in response to both variable affinities of Fabs to antigens (depending on maltose concentration) and of GA1CAR for Fab scaffolds (LRT (100 pM) vs. 4D5 (80 nM)). The functional capacity of GA1CAR-J cells was first measured by IL-2 release and CD69 expression using MBP coated wells (**Extended Data Fig 4A-B**). Activation levels were higher with soluble anti-MBP LRT Fab compared to the same Fab with an 4D5 scaffold (p <0.05), and insignificant with the non-binding Kappa Fab and untransduced cells (UTD). Using HEK cells expressing MBP as stimulators, IL-2 release was dose- dependent on Fab^LRT^ and Fab^4D5^ concentrations (**Extended Data Fig 4C-D**). Since the Fabs have identical affinities to the MBP target, the 15-fold difference in stimulation of IL- 2 secretion between these LRT and 4D5 Fabs is attributed to their differing affinities for the GA1 module. IL2 release by GA1CAR-J decreased with increasing maltose concentration (**Extended Data Fig 4E-F**), suggesting that lower binding affinity between the Fab and its target directly leads to decreased CAR-T activation.

The expression and functionality of GA1CAR were next evaluated in primary human CD8+ T cells. After lentiviral transduction, 40-98% of primary T cells achieved successful expression (**Fig. 2A**). In primary human CD8+ T cells transduced with GA1CAR (GA1CAR-T), IL-2 release was significantly higher with high-affinity Fab^LRT^ (EC50- 0.3 nM) compared to lower-affinity Fab^4D5^ (EC50 -4.8 nM) when targeting HEK- MBP cells (**Fig. 2B-C**). Similar results were observed for interferon-gamma (IFNγ) **(Fig. 2D)** and cytotoxicity **(Extended Data Fig 5)** when comparing high and low affinity Fab scaffolds. Notably, based on the EC50 values of the Fab^LRT^ scaffold, the stimulation of the IL-2 release was less sensitive than both IFNγ (4-fold) and cytotoxicity (20-fold) as a functional readout. Maximum responses were achieved using Fab concentrations in the 0.1- 10 nM range. Using different maltose concentrations, we demonstrated a Fab-to-target affinity-dependent release of IL-2 and IFNγ in GA1CAR-T cells (**Fig. 2E-G**). These data confirm the effective expression and regulatable function of GA1CAR on human primary CD8+ T cells and define the optimal range of Fab working conditions.

**Fig.2.**
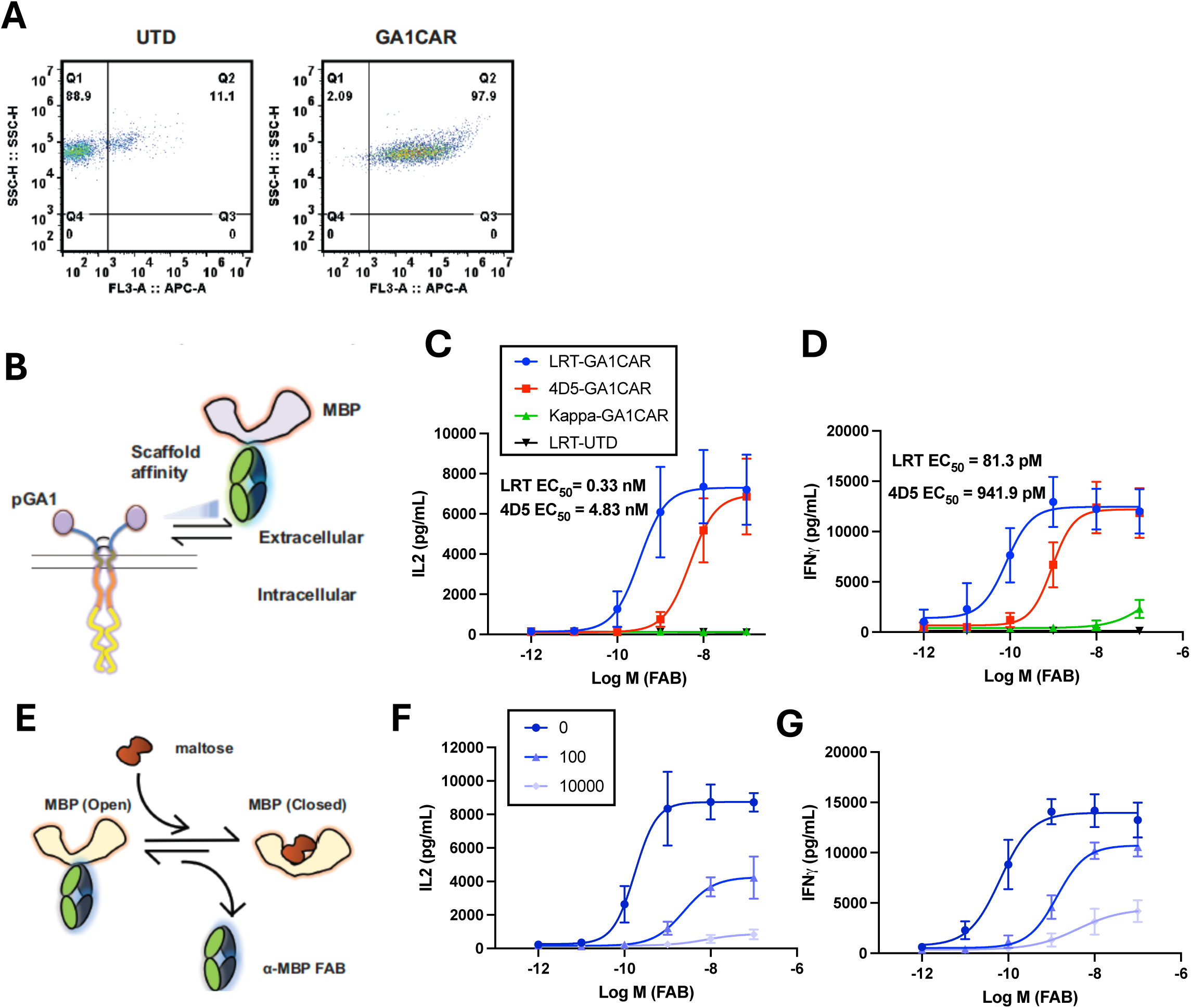
Tuneable function of GA1CAR expressed in primary human CD8^+^T cells. A) Surface expression of GA1CAR in human CD8^+^T cells, determined by flow cytometry by incubation with an irrelevant Fab^LRT^ followed by a secondary anti-human Fab antibody conjugated to Alexa Fluor® 647. B) Scheme for experiments determining the effect of using Fab scaffolds with varying affinities for the GA1CAR in GA1CAR-T cell function: C) IL2 release, D) IFNγ release. The data are presented as the mean ± SD, n = 3. E) Scheme for experiments determining the effect of varying the affinity of the Fab for its target in GA1CAR-T cell function, using the conformation-specific anti-MBP 70 Fab^LRT^ and HEK- MBP cells as targets. The affinity of the anti-MBP 70 Fab^LRT^ decreases with increasing maltose concentration. F) IL2, G) IFNγ release. The data are presented as the mean ± SD, n = 3.

### Cytotoxicity of GA1CAR Against Various Antigens and Cancer Cell Lines

To evaluate the GA1CAR system in a cancer-associated antigen model, the SKBR3 cell line, known for high HER2 expression (**Extended Data Fig 6A**), was used. Co-incubation of SKBR3 cells with GA1CAR-T cells loaded with different anti-HER2 Fab scaffolds resulted in IL-2 and IFNγ release and induced cytotoxicity when using the LRT or 4D5 scaffolds (p < 0.05), while only background levels were observed with the Kappa scaffold and anti-MBP-specific Fab **(Fig. 3A-C)**. Notably, scaffold affinity effects on T cell activation were observed only for IL-2 release (LRT > 4D5 > Kappa) but not for IFNγ or cytotoxicity, likely due to high antigen density masking these effects. Experiments with MDA-MB-231 cells with lower HER2 expression (**Extended Data Fig 6A**) showed that scaffold affinities influenced functional responses (**Extended Data Fig 6B-D**).

**Fig.3.**
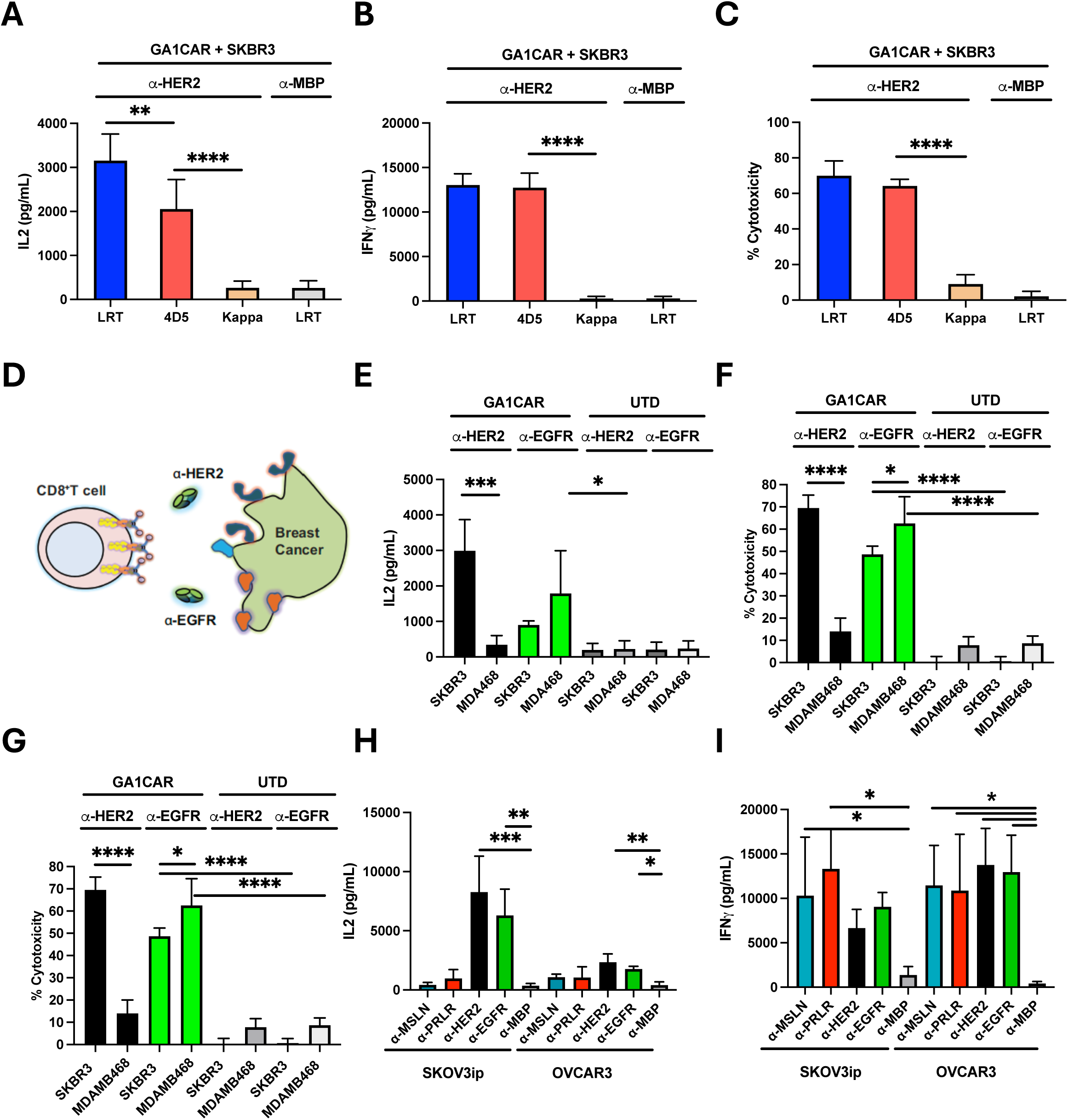
Recognition of multiple human cancer antigens by GA1CAR-T cells. A-C: GA1CAR-T cell function using anti-Her2 Fab scaffolds with different affinities for GA1CAR and SKBR3 cells as targets: A) IL2 release , B) IFNγ release, and C) Cellular cytotoxicity D) Cartoon depicting the recognition of SKBR3 or MDAMB468 breast cancer cell lines by GA1CAR, using either anti-HER2 or anti-EGFR Fab^LRT^ (1 nM). E) IL2 release, F) IFNγ release, G) Cellular cytotoxicity. H-I) Recognition of SKOV3ip or OVCAR3 ovarian cancer cell lines by GA1CAR, using either anti-HER2, EGFR, PRLR or MSLN Fab^LRT^ (1 nM). H) IL2 release, I) IFNγ release. Statistical significance was determined by one-way ANOVA followed by Šídák’s or Dunnett’s multiple comparison tests (*P < 0.05; **P < 0.01; ***P < 0.001; ****P < 0.0001), P < 0.05 considered significant. The data are presented as the mean ± SD, n = 3.

Further tests on breast cancer cell lines expressing EGFR and HER2 at different levels demonstrated that GA1CAR-T cells released IL-2 when anti-HER2 Fab^LRT^ was present with SKBR3 cells (p < 0.05), but not with MDAMB468 cells (**Fig. 3E)** due to undetectable HER2 levels (**Extended Data Fig 7A**). Anti-EGFR Fab^LRT^ induced IL-2 release in both SKBR3 and MDAMB468 cell lines, with similar trends for IFNγ release and cytotoxicity (**Fig. 3E-G**). Even low EGFR expression on SKBR3 (**Extended Data Fig 7A**) was sufficient for robust T-cell activation **(Fig. 3F-G).**

To explore additional cancer types, ovarian cancer cell lines SKOV3ip and OVCAR3 were used alongside Fabs targeting overexpressed receptors like MSLN, PRLR, HER2, and EGFR. GA1CAR-T cells released IL-2 **(Fig. 3H**) and IFNγ **(Fig. 3I)** in an antigen-specific way, even when some Fabs showed low flow cytometry staining levels (**Extended Data Fig 7B**). These results highlight GA1CAR’s versatility across different antigen-Fab^LRT^ pairs in breast and ovarian cancer cell lines.

### On-Off Switching Potential of Interchangeable Fab Scaffolds

A potential attribute of Fab scaffolds with varying affinities to the GA1 module is that they could be used to modulate GA1CAR activity through competitive displacement. To assess this potential, three Fab scaffolds with differing affinities—LRT (100 pM), SQLRT (10 nM), and 4D5 (50 nM)—were tested [27]. The activation of GA1CAR-T cells by SKBR3 cells through lower affinity (SQLRT or 4D5) Her2 Fab scaffolds was quenched adding a non-targeting Fab with higher affinity LRT scaffold as a competitive blocker (**Extended Data Fig 8A)**. The effects on IFNγ release and cytotoxicity were evaluated over time: 1 hr, 2 hrs, 4hrs and 8 hrs (**Extended Data Fig 8B-C)**). Results indicate that the competitive Fab^LRT^ reduced cytotoxicity in a time- and affinity-dependent manner; SQLRT showed little quenching at later time points for IFNγ release, suggesting rapid maximum activation post-CAR activation. These data demonstrate the potential to terminate GA1CAR-T cell function using a non-targeting competitive Fab with a higher affinity scaffold.

### In Vivo Efficacy of GA1CAR in Tumor Xenograft Mouse Models

Following the in vitro characterization of GA1CAR-T cells, their antitumor activity was evaluated using an in vivo tumor xenograft mouse model. Her2-expressing HCC1954 cells (5 million) were injected subcutaneously into the right flank of immunocompromised NSG mice (**Fig. 4A-B**). Once tumors were established, five mice per group were treated with GA1CAR-T cells, either pre-incubated with HER2-specific Fab^LRT^ or not, and conventional scFv HER2-CAR-T cells served as a control. Mice receiving GA1CAR-T and Fab^LRT^ received additional Fab^LRT^ injections every two days for two weeks. Results showed that GA1CAR-T cells combined with anti-HER2 Fab^LRT^ exhibited a robust antitumor effect (**Fig. 4B**). The activity of GA1CAR-T cells was comparable to that of conventional scFv CAR-T against HER2, while tumors in control groups without CAR-T cells or without Fab continued to grow.

**Fig.4.**
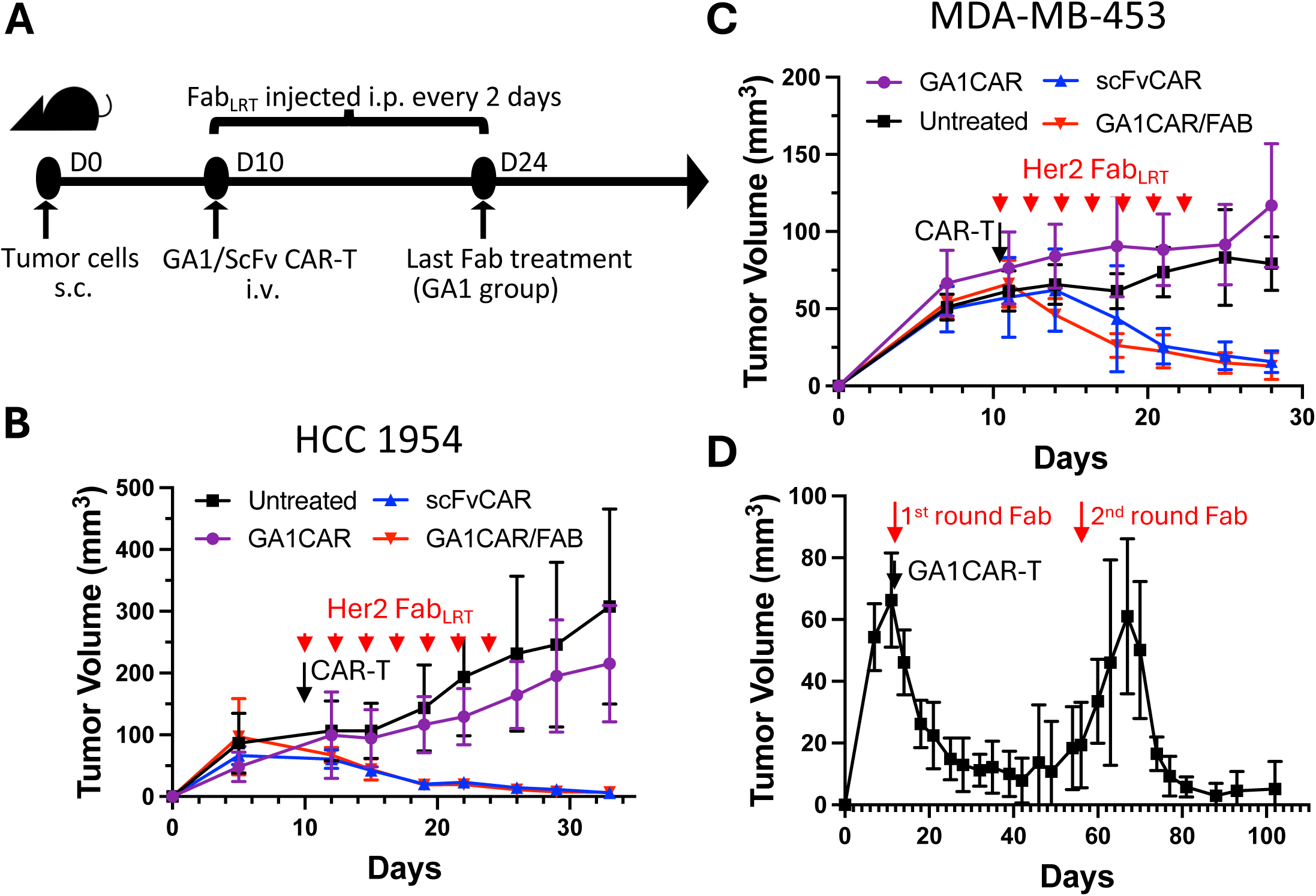
Anti-tumor activity of GA1CAR-T cells in breast cancer xenograft models. A) Experimental scheme. Female NSG mice were injected subcutaneously in the right flank with 5×10^6^ MDAMB453 or HCC1954 cells. Ten days later, the mice were injected with CAR-T cells i.v.. When indicated, Fab^LRT^ was preincubated with the GA1CAR-T cells for 30 min before GA1CAR-T injection and then injected i.p. every other day for 14 days at 200 ug/mouse. B-D) Tumor growth curves of NSG mice bearing HCC1954 (B) or MDAMB453 (C-D) tumors. Tumor-bearing NSG mice were treated with GA1CAR, Her2 scFvCAR , or GA1CAR/ Fab^LRT^ as indicated, or left untreated. Data are representative from two independent experiments for each cell line, with 5 animals per group. Statistical significance was determined by one-way repeated measures ANOVA followed by Dunnett’s multiple comparison tests. * indicates P < 0.05 for GA1CAR/Fab and scFv groups compared to GA1CAR control.

### Multiple Fab Dosing and Tumor Rebound

Although CAR-T therapy can reduce tumor mass to undetectable levels, residual tumor cells can persist, leading to nucleation and regrowth. The potential to "reactivate" GA1CAR by reintroducing Fabs was tested to address the rebound phenomenon.

MDAMB453 breast cancer cells were injected into NSG mice, using anti-HER2 Fab^LRT^ as a targeting agent. Initial injections began on day 10 and ended on day 24, resulting in rapid tumor contraction to baseline by day 27 (**Fig. 4C**). Tumor volume was assessed every three days and remained stable until day 50, after which it increased significantly, indicating rebound (**Fig. 4D**). On day 60, five additional anti-HER2 Fab^LRT^ injections were administered, leading to a rapid reduction in tumor volume. These findings suggest that GA1CAR remains active and potent for at least 100 days and can be reloaded with Fab^LRT^ for extended immunological therapy.

### Pharmacokinetics of GA1CAR-T Cells

To characterize the in vivo dynamics of GA1CAR-Fab^LRT^ coupling, experiments were conducted comparing two delivery methods: preincubating anti-HER2-Fab^LRT^ with GA1CAR-T cells before injection or delivering GA1CAR via retro-orbital injection followed by intraperitoneal HER2-Fab^LRT^ delivery. Notably, the HCC1954 tumor volume decreased similarly when comparing both delivery methods (**Extended Data Fig 9A**), indicating that the GA1CAR and Fab^LRT^ engage and couple efficiently in circulation upon injection. The duration of GA1-Fab^LRT^ interaction was assessed by measuring Fab+ GA1CAR-T cell percentages in peripheral blood post-Fab^LRT^ delivery. Eighteen hours after the first dose, comparable binding levels were observed regardless of preincubation (**Extended Data Fig 9B**). However, 48 hours after the third dose, no Fab^LRT^+ GA1CAR+ cells were detected, although GA1CAR+ cells persisted across all groups (**Extended Data Fig 9B- C**). This suggests that Fab^LRT^ binding duration is somewhat less than 48 hours and should be considered when designing dosing regimes.

The properties of CAR-T cells can be influenced by the PBMC donors from which they are derived [33], [34], [35]. To compare the effectiveness and kinetics of circulating GA1 vs scFv CAR-T cells, CD8+ T cells from three donors were transduced with either GA1CAR or scFv CARs, and their impact on tumor growth and circulating levels was compared using three recipient mice per CAR-T type and donor (**Extended Data Fig 10A**). Matched donor-GA1CAR-T/HER2 Fab^LRT^ and scFv CAR-Ts showed similar efficacy in controlling HCC1954 tumor growth, whereas tumors in mice treated only with GA1CAR or Fab^LRT^ grew progressively (**Extended Data Fig 10B**). Circulating CAR-T levels were more affected by donor origin than CAR type and decreased around tumor regression time for two out of three donor cohorts (**Extended Data Fig 10C**). CAR-T cell levels from Donor 1 increased after day 11 for both scFv and GA1CAR-T recipients, leading to adverse effects requiring euthanasia due to a potential graft-vs-host effect (ie non-tumor related). Overall, GA1CAR-T demonstrates similar antitumor efficacy and circulation kinetics as conventional scFv CAR-T cells, but offers enhanced targeting control through modulated Fab^LRT^ administration.

### Comparison of GA1CAR-T and scFv CAR-T Cell Phenotypes

In the context of CAR-T cell therapy, the expression of different CAR constructs can lead to distinct CAR-T cell phenotypes. The scFv CAR is known for low levels of downstream signaling due to tonic signaling, a result of CAR oligomerization caused by interactions between the scFv regions of neighboring CARs [36]. To explore how GA1CAR expression affects CAR-T cell phenotypes compared to scFv CAR, we conducted a bulk RNA sequencing analysis on FACS-sorted GA1CAR-T and scFv CAR- T cells isolated from tumors. In this study, matched GA1 or scFv CAR-T cells from two independent donors were transferred into groups of five NSG recipient mice each (**Fig. 5A**). On day six post-transfer, tumor infiltration by CAR-T cells was similar between GA1 and scFv recipients (**Fig. 5B, Extended Data Fig 11**).

**Fig. 5.**
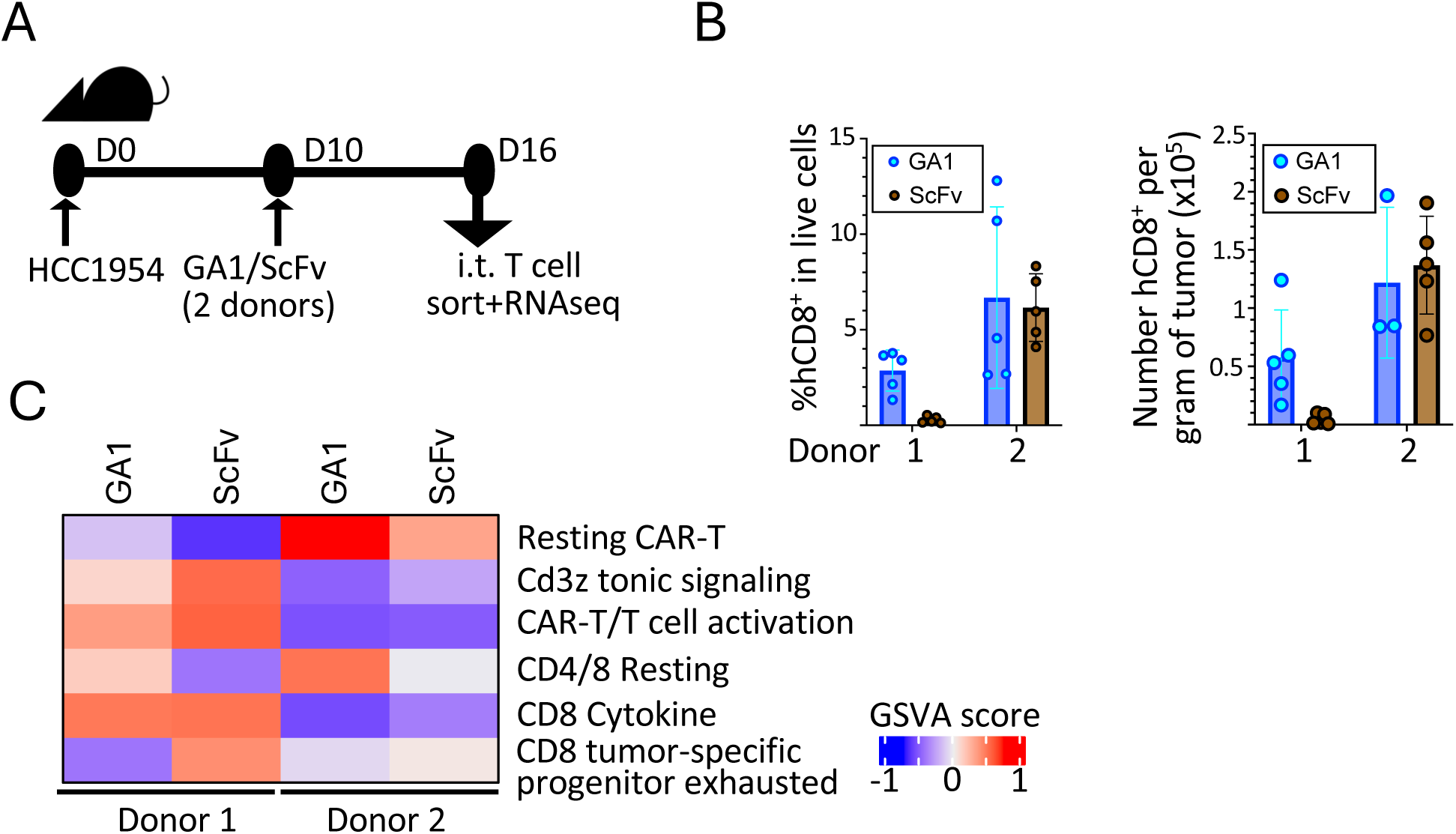
In vivo analysis of intratumoral GA1CAR-T cells vs. scFv CAR-T cells. A. Experimental scheme. Groups of 5 NSG mice were treated with donor-matched scFv/GA1CAR-T cells from two independent donors (4 groups total). B. Tumor-infiltrating CAR-T cells were quantified by flow cytometry at day 6 after CAR-T cell transfer. C. Enrichment in gene signatures indicative of resting/activated status was compared between FACS-sorted tumor-infiltrating scFv and GA1CAR-T cells by bulk RNAseq.

### RNA Sequencing and Phenotype Analysis

The RNA sequencing analysis focused on intratumoral T cells from two donors and two CAR-T types, examining gene signatures indicative of resting state, activation, or tonic signaling [37–40] (**Fig. 5C**). GA1CAR-T cells were enriched in resting CAR-T/T cell signatures for each donor. Conversely, scFv CAR-T cells showed enrichment in activation-associated signatures ("CAR-T CD3z tonic signaling” [38], "CD8 cytokine” [40], and "CD8 tumor-specific progenitor exhausted” [39].

To compare phenotypes in more controlled conditions, an *in vitro* activation model was used with donor-matched GA1 and scFv CAR-T cells from five independent donors (**Fig. 6A**). These cells were cultured in T cell media with maintenance cytokines (IL-2/IL- 15) for six days following CD3/CD28 bead removal. Subsequently, the CAR-T cells were co-cultured with HCC1954 HER2+ target cells (stimulation) or maintained in media (baseline) without cytokines for one day. GA1CAR-T cells at baseline were also cultured with and without Fab^LRT^ to assess whether Fab^LRT^ binding could trigger downstream ("tonic") signaling without antigen presence. FACS-sorted (**Extended Data Fig 12A**) CAR+ T cells from various conditions underwent bulk RNA sequencing (**Extended Data Fig 12B-C**) . Spectral flow cytometry was employed to profile the cells using markers associated with resting/memory or activation status at the protein level (**Fig. 6B-C**). Effector cytokine production (IFNγ, TNF) was measured in collected supernatants (**Fig. 6D**).

**Fig. 6.**
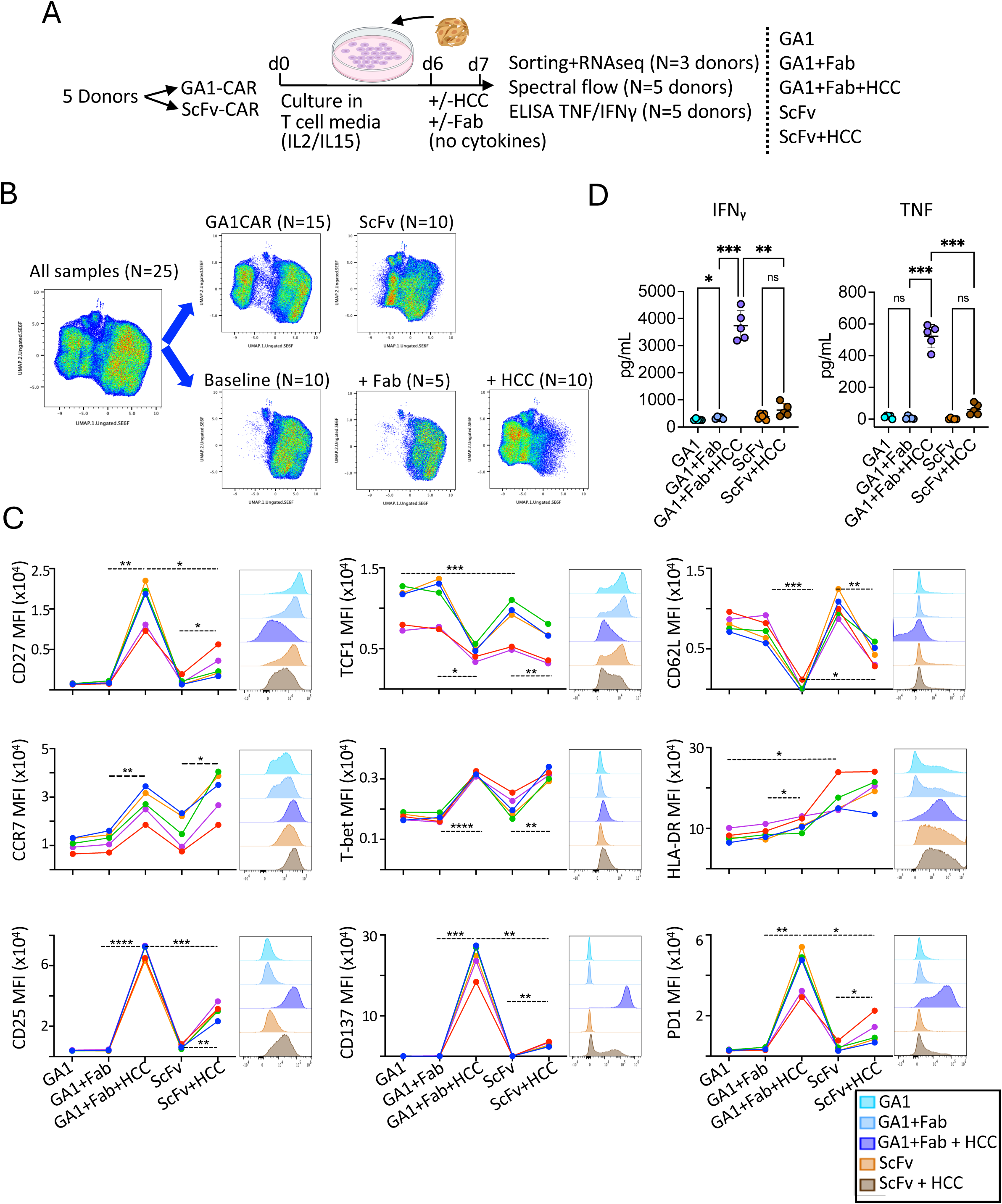
In vitro analysis of GA1CAR-T cells vs. scFv CAR-T cells. A. Experimental scheme. scFv and GA1/Fab^LRT^ CAR-T cells were generated from five independent donors. Equal numbers of CAR+ cells were activated in vitro with HCC1954 target cells and analyzed by RNAseq, spectral flow cytometry and ELISA on supernatants. B. UMAP distribution of samples analyzed by spectral flow cytometry. C. Expression of individual flow cytometry markers according to different experimental conditions compared. Each colored line represents a different donor. Representative histograms are shown for one donor. D. Levels of TNF and IFNg in supernatant. Each dot represents a different donor. Statistical significance in C-D was analyzed using repeated measurements ANOVA followed by Šídák’s multiple comparison tests (*P < 0.05; **P < 0.01; ***P < 0.001), ns:not significant. The data are presented as the mean ± SD, n = 5.

The gene expression analysis of CAR-T cells under baseline conditions highlighted a significant impact of adding Fab^LRT^ to GA1CAR-T cells. This addition was linked to an increased expression of CAR-T and T cell activation signatures (**Extended Data Fig 12B- C**). When comparing baseline GA1CAR-T and scFv CAR-T cells, the differences were subtle, but indicated a higher expression of genes related to T cell activation, like IFNγ, TNF, LAG3, and MHC-II, in resting scFv CAR-T cells compared to GA1CAR-T cells. Conversely, genes associated with resting or memory phenotypes, such as TCF7, LEF1, CD52, IL7R, and S100A4, showed higher expression in resting GA1CAR-T cells. Following stimulation with cancer cells, GA1CAR-T cells exhibited an overall increase in CAR-T and T cell activation signatures and a decrease in resting CAR-T and T cell signatures compared to scFv CAR-T (**Extended Data Fig 12B-C**). These patterns were generally corroborated at the protein level. Unsupervised UMAP clustering plots from spectral flow cytometry data demonstrated segregation based primarily on the samples’ activation status (**Fig. 6B**). The most notable differences were between baseline and cancer cell-stimulated GA1CAR-T samples; in contrast, baseline and stimulated scFv CAR-T samples were more similar. When individual marker expressions were analyzed (**Fig 6C**), minor differences were observed between GA1CAR-T and scFv CAR-T at baseline conditions. Statistically significant differences included higher TCF1 expression in GA1CAR-T (P=0.02) and higher HLA-DR expression in scFv CAR-T (P<0.0001). The addition of Fab^LRT^ to GA1CAR-T did not result in statistically significant protein-level changes. However, exposure to target cells led to substantial changes in protein expression, particularly an increase in proteins associated with T cell activation and effector function (CD25, CD137, PD1, T-bet) and a decrease in molecules linked to resting or memory states (TCF-1, CD62L, CD27). The changes induced by target cell exposure were often more pronounced for GA1CAR-T than for scFv CAR-T (CD137, PD- 1, TCF-1, CD27). For HLA-DR, the most significant differences between GA1 and scFv were at baseline; target cell incubation did not significantly increase expression. ELISA data (**Fig 6D**) indicated a significantly higher effector cytokine production for GA1CAR compared to scFv upon target cell stimulation (6-fold increase in IFNγ, P<0.0001; 8-fold increase in TNF, P<0.0001). This is likely due to the higher levels of surface CAR expression observed in GA1CAR-T compared to scFv CAR-T (**Extended Data Fig 13**). Collectively, these findings suggest that GA1CAR-T cells exhibit a more resting gene expression profile at baseline but become more activated upon direct antigen challenge and have a similar capacity for tumor infiltration as compared to scFv CAR-T cells.

### Redirecting GA1CAR-T Cells to new targets using Fab^LRT^ with different specificities

GA1CAR-T cells can be redirected to new targets using Fabs with different antigen specificities. The duration of Fabs binding and the effective lifetime of GA1CARs in circulation offer tunable control over their effector activity. This adaptability can be leveraged to alter the target specificity of the CAR-T cells in a "plug-and-play" manner. To test the feasibility of this approach, a model was developed using NSG mice bearing HCC1954 tumors (characterized by low EGFR and high HER2 levels) treated with GA1CAR-T cells alongside anti-EGFR Fab^LRT^ for ten days (**Fig. 7**). Targeting EGFR significantly slowed tumor growth but did not eradicate HCC1954 tumors due to the low EGFR levels in this cell line. On day 25, half of the mice received a second round of Fab^LRT^ doses targeting HER2. Switching Fabs to target a second antigen allowed GA1CARs to regain control over tumor growth, leading to extended survival for the mice (P=0.0007). These results demonstrate the potential of using GA1CARs for targeting multiple antigens following a single infusion of CAR-T cells by adjusting the specificity of the Fabs as needed.

**Fig. 7.**
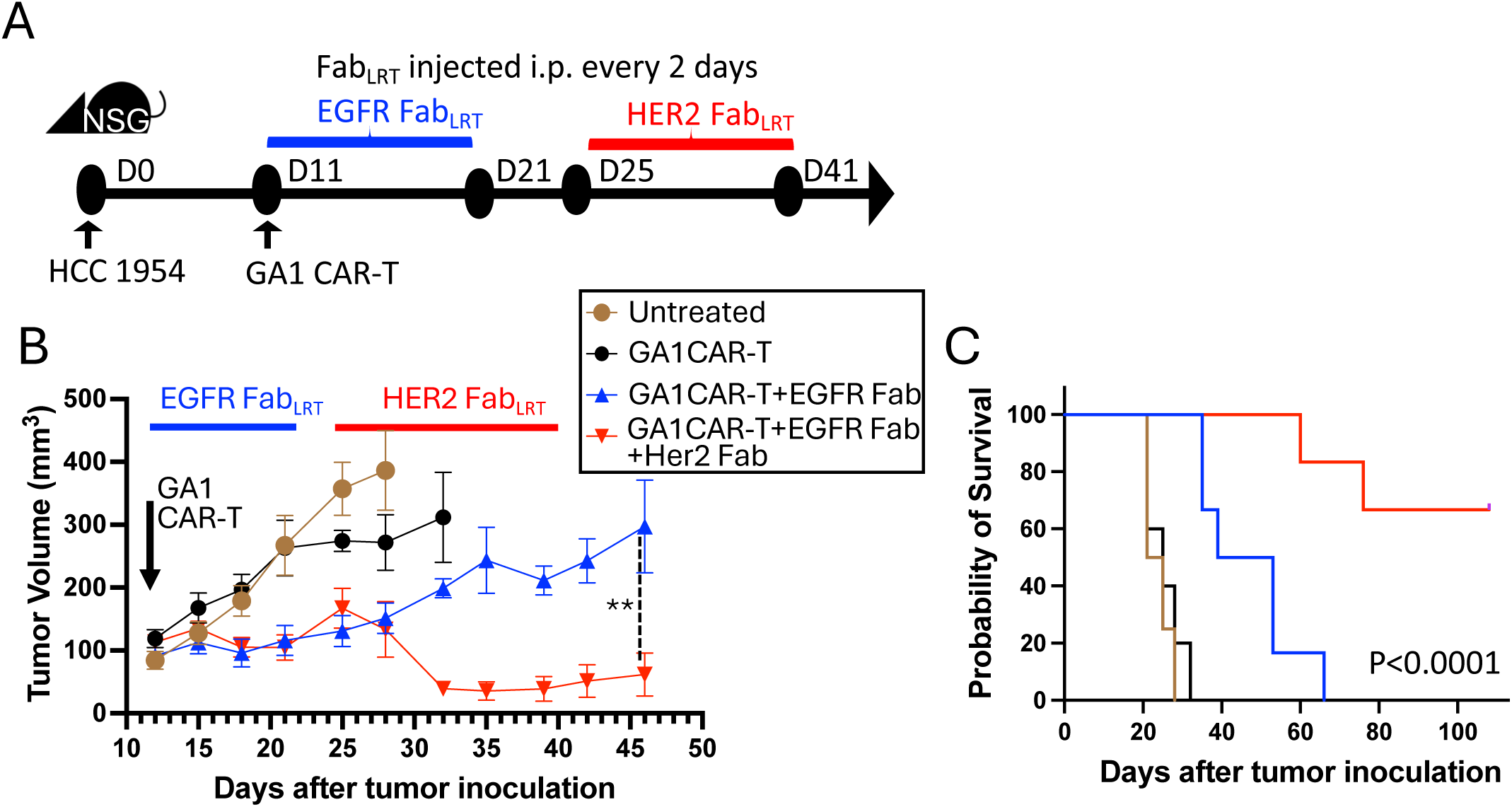
Switching Fabs to redirect GA1CAR-T cells to a new target in vivo. A. Experimental scheme. HCC1954 tumor-bearing mice received GA1CAR-T cells with or without treatment with EGFR Fab^LRT^ for 10 days. After that, half of the EGFR Fab^LRT^-treated animals received a second round of Fab^LRT^ for 16 days, this time specific for HER2, whereas the other half received no further treatment. N=5 for control groups (untreated, GA1CAR only) N=6 Fab-receiving groups (EGFR only, EGFR+HER2). B. Tumor volumes per group (average+/SEM). Data was analyzed using repeated measurements ANOVA followed by Dunnett’s multiple comparison tests (*P < 0.05; **P < 0.01; ***P < 0.001; ****P < 0.0001), P < 0.05 considered significant. C. Kaplan-Meier survival plot was analyzed with log-rank (Mantel-Cox) test .

## Discussion

This study presents a comprehensive evaluation of the GA1CAR system, a modular chimeric antigen receptor designed to enhance the specificity and adaptability of CAR-T cell therapies. The GA1CAR design addresses a number of the technical challenges that are faced by traditional CAR-T systems; in particular potential toxicity, target heterogeneity, the emergence of antigen-loss variants, and the complexity of manufacturing [41]. Its design is simpler than most existing universal CARs, with the targeting component—a Fab—being easier to manufacture than a full CAR-T product. The results demonstrate the efficacy and versatility of GA1CAR in various *in vitro* and *in vivo* settings, highlighting its potential for clinical applications.

The GA1CAR-T system is modular and tunable, allowing for the interchange of its targeting moiety in a "plug-and-play" fashion based on tumor antigen expression profiles. This system offers control over potency and signaling duration to minimize adverse immunological responses. Additionally, it is tag-less, making it less susceptible to cleavage by proteases. Our proof-of-concept studies utilized a second-generation CAR construct; however, the GA1CAR attributes can be easily integrated into next-generation constructs to leverage ongoing advances in CAR technology.

The GA1CAR system is built on an engineered protein G variant (GA1), which binds effectively to various engineered Fab scaffolds and is non-immunogenic [27], [29]. These scaffolds differ by only a few strategically placed mutations, yet produce differences in affinities ranging across three orders of magnitude, allowing for broad manipulation of engagement [27–28]. The engineered Fabs function similarly to single-molecule Bispecific T-cell Engagers (BiTEs), binding tightly and specifically to both the target antigen on cancer cells and the CAR through its GA1 moiety. This dual modulation—by varying Fab affinities or using different affinity scaffolds—offers significant control over treatment and toxicity management. For example, a lower-affinity scaffold can be used initially and replaced with a higher-affinity scaffold, if well tolerated. To characterize GA1CAR as a versatile universal system capable of targeting multiple antigens on cancer cells with high selectivity and efficacy *in vivo*, we used immortalized and primary human CD8+ T cells alongside breast and ovarian cancer cell lines expressing various endogenous surface antigens. *In vivo* experiments demonstrated GA1CAR-T cells’ effectiveness in two breast cancer models targeting two different antigens (Her2 and EGFR). Sequential administration of Fab^LRT^ with different specificities improved outcomes when initial treatments failed to fully control tumor growth. Further, our results demonstrate that GA1CAR can address tumor escape caused by heterogeneity or low antigen expression by targeting multiple antigens.

Although recent advances in regulatable CAR formats offer high control levels [24–26], the conventional scFv CAR format remains the most studied and serves as the standard for comparison. In our mouse xenograft model, GA1CAR showed comparable efficacy in reducing tumors when compared to conventional scFv CARs. While scFv CARs are always loaded due to their covalent attachment to their targeting moiety— leading to potential toxic side effects—the GA1CAR format offers several safety advantages. The Fab^LRT^s used with GA1CAR have a finite lifetime in peripheral blood (∼2 days), but can be reloaded as needed through additional dosing. This flexibility contrasts with scFv CARs’ rigidity and associated safety risks from on-target/off-tumor toxicities due to binding similar antigens in healthy tissues. Another key advantage of the GA1CAR system is its ability to target multiple tumor antigens simultaneously or sequentially, reducing the risk of antigen-negative escape. Unlike other systems that use small peptide tags prone to cleavage by proteases, GA1CAR relies on stable protein-protein interactions between GA1 and Fab^LRT^, minimizing susceptibility to cleavage. GA1CAR’s design is also simpler and less structurally constrained than other modular CAR systems.

When comparing transduction rates and expression levels between GA1CAR and scFv CAR constructs generated in the same lentiviral vector, GA1CAR consistently showed higher transduction rates and expression levels in recipient T cells. At baseline, gene expression data suggested that GA1CAR-T cells were more quiescent than scFv CAR-T cells in the absence of Fab^LRT^; however, adding Fab^LRT^ partially triggered activation signatures without increasing cytokine production or altering surface marker expression. This suggests that if tonic signaling occurs, it has limited consequences at the protein level. Importantly, our universal GA1CAR only recognizes Fabs with LRT or 4D5 mutations—reducing the risk of autoimmune reactions from cross-reactivity with other human antibodies. We anticipate the system’s design will facilitate IgG formats as targeting molecules. While Fabs offer precise control over dose and activity with better tissue penetration, IgGs provide benefits such as bivalency and longer serum half-life due to FcRn-mediated recycling. Future studies will explore using IgGs with GA1CAR-T cells and further optimize hinge, transmembrane, and intracellular domains for specific antigen targets [42]. Specific spatial orientations between T cells and target cells are crucial for efficient immunological synapse formation—a factor that will be considered in future optimization efforts. Additionally, new strategies could enhance effector functions or safety mechanisms for GA1CAR-T cells through co-expression of cytokines or homing receptors or through suicide genes and intracellular split domains. Non-viral GA1CAR delivery methods may also be explored [43–45].

In summary, we have developed an orthogonal universal CAR-T system capable of sequential or simultaneous targeting of multiple cancer antigens while offering enhanced stability, control, and delivery advantages compared to conventional systems. The GA1CAR system can also serve as a preclinical platform for screening new Fab^LRT^ molecules from high-throughput antibody discovery pipelines for personalized medicine applications in adoptive cell therapy.

### Materials and Methods Cell lines

HEK293T, Jurkat, SKBR3, MDAMB231, MDAMB468, MDAMB453, and HCC1954 Cell lines were purchased from ATCC. HEK-Flp cells were from Invitrogen. The ovarian cancer cells SKOV3ip and OVCAR3 were provided by Dr. Ernst Lengyel at the University of Chicago. Most cells were grown in RPMI media with 10% heat inactivated FBS, 100 U/mL of penicillin, and 100 mg/mL of streptomycin. HEK293T cells were grown in DMEM media with 10% heat inactivated FBS, 100 U/mL of penicillin, and 100 mg/mL of streptomycin.

### HEK-MBP stable cell

The HEK-MBP stable cell line was created using the HEK Flp-in system following the vendor protocol (Invitrogen K6010-01). Briefly, a chimeric receptor composed of a mouse IgG kappa signal peptide, full MBP as ectodomain, PDGFR transmembrane, and intracellular GFP, was cloned into pcDNA^TM^ 5/FRT expression vector using XhoI/XbaI cloning sites. The plasmid expressing this chimera (pcDNA^TM^ 5-MBP/FRT, for simplicity) and the plasmid pOG44 (expressing the Flp recombinase) were electroporated into HEK Flp-in cell line. Stable clones selected with hygromacyn B at 100µg/mL, and single clones were confirmed by flow cytometry with anti-MBP Fabs and intracellular GFP expression.

### PBMC and CD8^+^T cell isolation

Peripheral blood mononuclear cells (PBMC) were isolated from leukopak reduction filters obtained by apheresis of de-identified healthy donors from the Blood bank at the University of Chicago Medicine, via density gradient centrifugation using Ficoll (Corning, 25-072-CV). CD8^+^T cells were purified using the commercial kit from Miltenyi Biotech, following the instructions suggested. Purity of the separation was monitored by flow cytometry following the manufacturer recommendation. CD8^+^T were frozen in 50% FBS, 40% RPMI, 10% DMSO at a density of 50-60 ×10^6^/vial. The final yield of CD8^+^T ranged from 20-60×10^6^ cells per reduction filter

### Lentivirus vector and packaging

The sequence of protein GA1 variant -with a K32E mutation to reduce binding to human Fc [29]. (TPAVTTYKLVINGRTLSGYTTTTAVDAA<u>TAE**K**</u>VFKQYAYVHEVDGEWTYDDATKTFTV TEKPEKL) - was cloned into a pLVX vector (Takara, 631987) via Infusion cloning (Takara, 638944). GA1 sequence was cloned in-frame and upstream of a CD8 hinge, CD28 transmembrane, a 41BB co-stimulatory domain, and CD3ζ domain (full amino acid sequence in Extended Data Fig 1). Lentivirus particles were packaged in HEK293T cells plated in four T-225 flasks in HEK media (10% FBS, DMEM, 1 penicillin 100 U/mL/ streptomycin 100 µg/mL). The lentivirus vector was co-transfected with psPAX2 (Addgene 12260) and pMD2.G (addgene 12259) using lipofectamine LTX reagent (ThermoFisher, 15338500) at a ratio of 1:0.9:0.21, respectively, with 228 µg total DNA. Fresh media was added after 18 hr and lentivirus particles harvested at 48 hr and 72 hr after transfection. Lentivirus was concentrated 10-fold via precipitation with lenti-X concentrator (Takara, 631232) and re-suspended in PBMC media (RPMI media containing 10% FBS, 2x L-Glutamine, 1mM sodium pyruvate, 1x non-essential amino acids and vitamins, and penicillin 100 U/mL/ streptomycin 100 µg/mL) before aliquoting and freezing. Viral titer was monitored by Lenti-X GoStix Plus (Takara, 631280) and quantified by qPCR (Takara, 631235).

### Lentivirus transduction

50×10^6^ human CD8^+^T cells were plated in a T-175 flask and activated with Dynabeads Human T-Activator CD3/CD28 (ThermoFisher, Cat.No. 111.31D) for 24 hours, following the manufacturer recommendations. After 24 hr of activation, cells were transferred into a 6-well plate at 8 ×10^6^ cells/well and transduced with 3.5 ml of concentrated lentivirus particles by spinoculation at 1000 g for 2 hr at 32°C in media containing IL-2 at 50 U/µL and polybrene at 6 µg/mL. After transduction, cells were incubated overnight, and fresh media added next morning containing 50 U/mL IL-2 and 50ng/mL IL-15. Cells were grown for up to two weeks before frozen in 50% FBS, 40% RPMI, 10% DMSO at a density of 50-60 ×10^6^/vial.

### Cytokine release assay

For cytokine release by GA1CAR transduced in Jurkat cells, ELISA plates (Greiner bio one, 655081) were coated for 2 hr with neutravidin (2µg/mL) and 1 hr with MBP-biotin (100nM). Next, 10×10^5^ GA1CAR cells per well were incubated with Fab at 10nM overnight. Additionally, 10×10^5^ GA1CAR cells were incubated with 1×10^5^ HEK-MBP target cells and different Fab scaffolds and concentrations. Cells were co-cultured overnight in a 96-well U-bottom shape plate (Greiner, 650261) in fresh media without cytokines. Next day, supernatants were collected and analyzed for human IFN γ and IL2 using commercial kits from Cisbio. (Cat No. 62HIFNGPEH and Cat No. 62HIL02PEH, respectively). In the case of human CD8^+^T cells, 10×10^5^ CD8T cells transduced with GA1CAR and 1×10^5^ target cells were co-cultured overnight in a 96-well U-bottom shape plate in fresh media without cytokines. Next day, supernatants were collected and analyzed for human IFN γ and IL2 using commercial kits from Cisbio.

### Cytotoxicity assay

Cytotoxicity was measured by the release of lactate dehydrogenase (LDH) into the cultured media. Briefly, effector and target cell at 10:1 ratio were incubated with different Fabs concentrations for 16 hrs. Cytotoxicity of CAR-T cells toward target cells was measured using the non-radioactive cytotoxicity assay CytoTox-96 ® from Promega (Cat. No. G1780) as per manufacturer’s protocol.

### scFv/GA1CAR-T cell activation in vitro

CD8+ T cells were isolated from five healthy donors and subsequently used to generate donor-matched GA1 and scFv CAR-T cells. Following CD3/CD28 bead activation and transduction, CAR-T cells were expanded in RPMI medium supplemented with 10% fetal bovine serum (FBS), 1% penicillin-streptomycin (P/S), 2 mM L-glutamine, 1 mM sodium pyruvate, non-essential amino acids (NEAA, 1X), 50 U/ml IL-2, and 50 ng/ml IL-15 for 10 days post-transduction. On day 10, human T cell activator CD3/CD28 beads were removed, and cells were washed twice with cytokine-free medium. GA1CAR-T cells were then incubated with or without Her2 Fab^LRT^ for 30 minutes in cytokine-free medium. GA1CAR-T cells incubated or not with Her2 Fab^LRT^ (200 nM) and Her2-specific scFv CAR-T cells were co-cultured with HCC1954 cells in duplicates, in 12-well tissue culture plates in a volume of 1.5 ml per well for 24 hours at 37 °C in a 5% CO2 incubator. The CAR-T to target cells ratio was 3:1 (5 × 10⁶ CAR-T cells to 1.6 × 10⁶ target cells). After 24 hours, cell supernatants were collected and stored at -80°C for subsequent cytokine analysis using Biolegend ELISA Kits: TNF-α (cat # 430204) and IFN-γ (cat # 430104).

Half of the cells from each sample were allocated for single-cell RNA sequencing (scRNA-seq) after extraction of RNA using RNeasy Mini Kit (Qiagen), and the remaining half was used for immunophenotyping using flow cytometry.

### Intratumoral scFv/GA1CAR-T cell analysis ex vivo

Donor-matched GA1 and scFv CAR-T cells were injected i.v. into HCC1954-tumor bearing NSG mice at day 12 after tumor inoculation, at 4 million CAR^+^ cells/mouse. Two different donors were used and 5 recipient mice/group, for a total of 4 groups and 20 mice. GA1-recipient mice were treated with 3 doses of Her2 Fab^LRT^ i.p. (200 ug/dose) at days 0, 2 and 4 since GA1CAR-T cell transfer. At day 6, all mice were sacrificed. Tumors were excised, weighed, cut into 1 mm fragments and digested for 25 minutes at 37C with 75 ug/mL Liberase TM (Roche) and 20 ug/mL DNase I (Sigma) under gentle shaking. Cell suspensions were filtered using 70 uM cell strainers. Half of each individual mouse sample was used for quantification of infiltrating CD8+T cells/g of tumor by flow cytometry by staining with Live/Dead Yellow Dead Cell Stain (Invitrogen) and anti-human CD8 APC (Biolegend) and acquisition using a Cytek Aurora flow cytometer. The other half of each individual tumor sample was pooled per group (ie cells from the 5 mice from each group were pooled into a single sample) before magnetic enrichment for human CD8^+^ T cells using CD8 MicroBeads from Miltenyi (130-045-201). Pooled CD8-enriched samples were then stained with Live/Dead Yellow Dead Cell Stain and anti-human CD8 APC. CD8^+^Live/Dead^-^ cells were sorted in a FACS Aria into tubes containing plain RPMI. Pellets were processed and RNA was extracted according to the instructions of the RNeasy Micro Kit (Qiagen).

### Flow cytometry

Cell staining in figure 1 was done using different Fab concentrations in cell media (10% FBS, RPMI, P/S) at 37°C for 30 min, followed by two washes, and secondary antibody Alexa-647 staining (Jackson immunoresearch, catalog 109-605-006) at 400x dilution and at 4°C. CD69 expression in figure 2 was measured by flow cytometry using an anti-CD69 antibody conjugated to APC (Biolegend, 310910). Briefly, 0.1-0.2×10^6^ cells were stained in 50 µl of 1:40 antibody dilution in 1xPBS buffer, incubated for 30 min, washed twice with 1xPBS, and fixed in 0.5% PFA before flow cytometry analysis.

For ex vivo and in vitro GA1 and scFv immunophenotyping: cells were washed with PBS before staining with Live/Dead Blue Fixable dye ( # L34961A, Invitrogen) for 30 min at RT, washed with PBS, stained with surface antibodies for 15-30 min at 4C, washed again and fixed using Cytofix (cat# 554655, BD Bioscience). GA1CAR-T samples were preincubated with an irrelevant Fab^LRT^ at 200 nM for 30 min before staining with an anti-F(ab’)2 antibody (details below) for detection of the GA1 CAR. Cells were kept in flow buffer until analysis. For intracellular staining, the Foxp3/Transcription Factor Staining Buffer Set (cat # 00-5521-00) from Invitrogen was used according to the manufacturer’s instructions. The following anti-human antibodies were used for CAR-T cell immunophenotyping: anti-CCR7-BV421 (cat # 353208), anti-CD62L-BV605 (cat # 304833), anti-CD127-BV650 (cat # 351325), anti-CD25-BV711 (cat # 302635), anti-Fas- BV785 (cat # 305645), ant-CD27-FITC (cat # 302806), anti- CD52-PreCP-Cy5.5 (318911), anti-TIM3-PE (cat# 345006), anti-CD137-PE-Daazzle 594 (cat # 309825), anti-HLA-DR-PE CY7(cat # 307607), anti-PD1-PE-CY7 (cat # 329917), anti-CD39-APC Fire 750 (cat # 328229), anti-Ki67-BV711 ( cat # 350515), anti-Tbet-FITC (cat # 644811) from Biolegend; anti-TCF1/TCF7-PE ( cat # 14456S) and anti-LEF1-AF700 (cat # 38921S) from Cell signalling; anti-F(ab’)2-AF647 (cat # 109-606-097) from Jackson immunoresearch laboratories; and anti-CD8-BUV805 (cat. # 612889) from BD Bioscience. NSG mouse blood samples were surface-stained, treated with NH4Cl red blood cell lysis buffer and immediately acquired (without washing) after addition of CountBright beads (Invitrogen) to determine the absolute counts of CAR-T cells in peripheral blood. Samples were analyzed using a Cytek Aurora spectral flow cytometer (ex vivo/in vitro phenotyping) or BD Fortessa (circulating CAR-T analysis) at the University of Chicago Cytometry and Antibody Technology Core. Flow cytometry data was analyzed using FlowJo v10.10 software (FlowJo LLC, BD Life Sciences, Ashland, OR, USA)

### Single Molecule Pulldown Assay, Spot Counting, Photobleaching Analysis

Previously described methods were followed [30], [46]. In brief, quartz slides were passivated with 200 mg/mL methoxy polyethylene glycol (PEG) containing 2.5% (wt/wt) biotinylated PEG. HEK293 cells were transfected with CAR constructs (1.5×10^6^ cells in 6-well plate) and lysed after 48 hours in 200 µl of lysis buffer (50 mM Tris pH=7.4, 120mM NaCl, 1% NP-40, 5mM EDTA, 25 mM NaF, 25 mM Sodium pyrophosphate, 25mM β - glycerophosphate, 1mM PMSF, 1 ug/mL Leupeptin, 1 ug/mL chymostatin, 1 ug/mL pepstatin, 1 ug/mL antipain). Cell lysate was cleared by centrifugation at 10,000 x g for 10 min, diluted 100-fold or enough to obtain a surface density optimal for single-molecule analysis. For SNAP tag labeling, 100 µl of non-diluted lysate was labeled with 1 uL of 0.6 mM dye, incubated for 30 min at 37°C, and pass through a spin column (PD MiniTrap G- 10, Cytiva) to remove free dye, fractions were collected and used for single molecule pulldown. Mean spot count per image and SD were calculated from images taken from 10 or more different regions. For photobleaching analysis, 500 frame videos were recorded using a 200 ms exposure, and single molecule fluorescent time traces of surface immobilized proteins were manually scored as having one to four bleaching steps or were discarded if no clean bleaching steps could be identified. Total number of molecules successfully scored as bleaching in one to four steps (N) is depicted in the figures. Single-molecule data was collected in a custom-built TIRF microscope equipped with a 100X 1.40 NA objective (Olympus) and EMCCD camera (iXon, Andor technologies). All experiments were performed at room temperature (22-25°C).

### Analysis of gene expression

RNA samples were analyzed by the The University of Chicago Functional Genomics Facility (RRID:SCR_019196). RNA quality and quantity was assessed using the Agilent bio-analyzer. Strand-specific RNA-SEQ libraries were prepared using a TruSEQ mRNA-SEQ library preparation protocol (Illumina provided). Library quality and quantity was assessed using the Agilent bio-analyzer and libraries were sequenced using an Illumina NovaSEQ6000 (Illumina provided reagents and protocols). RNAseq data was preprocessed using the nf-core ^60^ rnaseq v3.14.0 pipeline (doi: https://doi.org/10.5281/zenodo.1400710). Briefly, read QC was performed using FastQC v8.30 , read alignment using STAR v2.7.9a, duplicate marking using Picard v3.0.0, and read quantification using Salmon v1.10.1. The pipeline was executed with the following command: nextflow run nf-core/rnaseq -r 3.14.0 -profile singularity --input sample_info.csv --max_memory 48.GB --genome GRCh38G34 --gencode --skip_preseq false --skip_rseqc true --pseudo_aligner salmon --outdir results -resume. Raw reads were then converted to counts-per-million, filtered for low gene expression, and normalized by library size using trimmed mean of M-values normalization in R v4.2.1 using the EdgeR v.3.38.4 package.

### Fab cloning

Fab against MBP have been previously reported by our lab [32]. Fab against HER2 was cloned as described previously [28]. Fabs against PRLR were generated by phage display selection using recombinant prolactin receptor fused to Fc fragment. Briefly, the humanized CDR-containing regions were cloned into pSFV4 or pRH2.2 vectors, via Infusion cloning. Fabs against EGFR and MSLN were cloned using the VH and VL chains sequences published in #US 6235883B1 and #WO2015090230A1 patents, respectively. All VH and VL sequences (including LRT mutations), were fused upstream of CH and CL chains, and were chemically synthesized, as gBlocks by IDT. The gBlocks were inserted into pRH2.2 vector via infusion cloning, for protein expression. For IgG cloning, the variable region from the light was PCR with forward primer (5- TTACGTTCGTCGCGGCCGCAGTCGCCTCCGATATCCAGATGACCCAGTC-3) and reverse primer (5-GCCAAGCTGGGGATCCTTACTTACTTAGCTGCTGCTCAACACTCTCC CCTGTTGAAGCTC-3). The PCR product was cloned into a pSCSTa vector digested with NotI and BamHI enzymes. The heavy chain was PCR with forward primer (5-TTACGTTCGTCGCGGCCGCAGTCGCCGAGATCTCCGAGGTTCAGCTG-3) and reverse primer (5-TGGGCCCTTGGTGCTAGCCGAGGAGACGGTGACCAGGGTTC-3).

The PCR product was cloned into a pSCSTa vector containing the human IgG Fc domain, digested with NotI and NheI enzymes. Cloning was done via InFusion cloning (Takara, 638944) and positive cloning was verified by DNA sequencing.

### Fab purification

Fabs were cloned in pRH2.2 or pSFV4 vectors (IPTG inducible) for bacterial expression. Fabs were expressed in the periplasm of *E. coli* BL21 (Gold). Cells were grown in 2xYT medium for 4-5 hr at 37°C. After induction by 1mM IPTG at 0.8-1.2 OD600, cells were harvested by centrifugation and sonicated in 50mM Tris, pH =7.5, 200 mM NaCl, 2mM PMSF. The centrifugation cleared lysates were heated at 60°C for 30 min. Next, the lysate was centrifuged 40 min at 4000g to removed precipitated proteins. Lysates were filtered through a 0.2 µm filter and purified on AKTA purifier equipped with a home-made Protein GA1 resin (SulfoLink Coupling Resin, Thermo Scientific). Next, peak fractions were loaded onto an ion exchange Resource S 1-ml column (GE Healthcare). After washing with 50 mM sodium acetate (pH 5.0), Fabs were eluted with a linear 0%–100% gradient of buffer containing 50 mM sodium acetate (pH 5.0) and 2 M sodium chloride. Fractions containing pure Fab were pooled, neutralized with 50 mM HEPES (pH 7.5), dialyzed against buffer containing 50 mM HEPES (pH 7.5) and 200 mM sodium chloride or 1xPBS, concentrated, and stored in aliquots at -80°C.

### Endotoxin removal and quantification

Endotoxin removal from Fabs was done with Proteus NoEndo columns (GEN- NoE24Micro; Protein Ark), and final levels quantified using the Limulus Amebocyte Lysate assay (88282; Thermo Fisher Scientific). The final endotoxin levels in the Fab preps were below 5 EU/mg.

### Xenograft mouse model

Animal care and use were in accordance with institutional and NIH protocols and guidelines. All studies were approved by the Animal Care and Use Committee of The University of Chicago. All mice were maintained under specific pathogen-free conditions. Six to eight-week-old female and male NSG (NOD. Cg-Prkdc^scid^ Il2rg^tm1Wjl^/SzJ) mice were purchased from The Jackson Laboratory. To initiate the studies, 3-5×10^6^ HCC1954 or MDAMB453 cells in 50% growth factor reduced-matrigel (BD Biosciences, cat. 356231) were injected subcutaneously into the right flank of mice. Once tumor reached a volume of 100-150 mm^3^ , mice received 10-20×10^6^ of CAR+ scFv or GA1CAR-T cells by retro-orbital injection preincubated or not with α-HER2 Fab^LRT^ for 30 minutes, followed by a dose of anti-HER2 Fab^LRT^ every other day for 10-14 days. In some experiments, mice received anti-EGFR Fab^LRT^. All Fab^LRT^s (in 1xPBS buffer) were injected intraperitoneally at 200 µg/mouse. In control groups, mice were injected with GA1CAR-T cells only or received no treatment. Mice were monitored and tumor was measured with a caliper every 3-4 days, and tumor volume calculated by WxHxL/2. In HCC194-bearing mice, tumors in untreated control groups often experience spontaneous ulceration and /or leakage of fluid when they reach 300-500 mm^3^ volume and need to be euthanized, therefore we used a 300 mm^3^ tumor volume endpoint for the survival plot in Fig 8C. Peripheral blood was obtained from the retro-orbital sinus under general anesthesia.

## Statistical Analysis

All experimental data are presented as mean ± SD, of at least three sets of independent experiments, unless otherwise indicated. Data analysis was performed using GraphPad Prism software (GraphPad Software, San Diego, CA).

## Acknowledgments

We thank all Kossiakoff and Weichselbaum lab members for valuable suggestions and assistance with experiments and Mark Eckert (Ernst Lengyel lab) for providing the ovarian cell lines.

## Author contributions

A.A.K. conceived the project; A.A., E.A., K.W., E.M., A.S., L.W., T.S., A.Z. and K.D. designed and performed the experiments; Z.P.S., S.U., and M.L. generated new and critical reagents, A.A., E.A., R.R.W. and A.A.K. wrote the paper. All authors reviewed and approved the final manuscript.

---

**Extended Data Fig 1.**
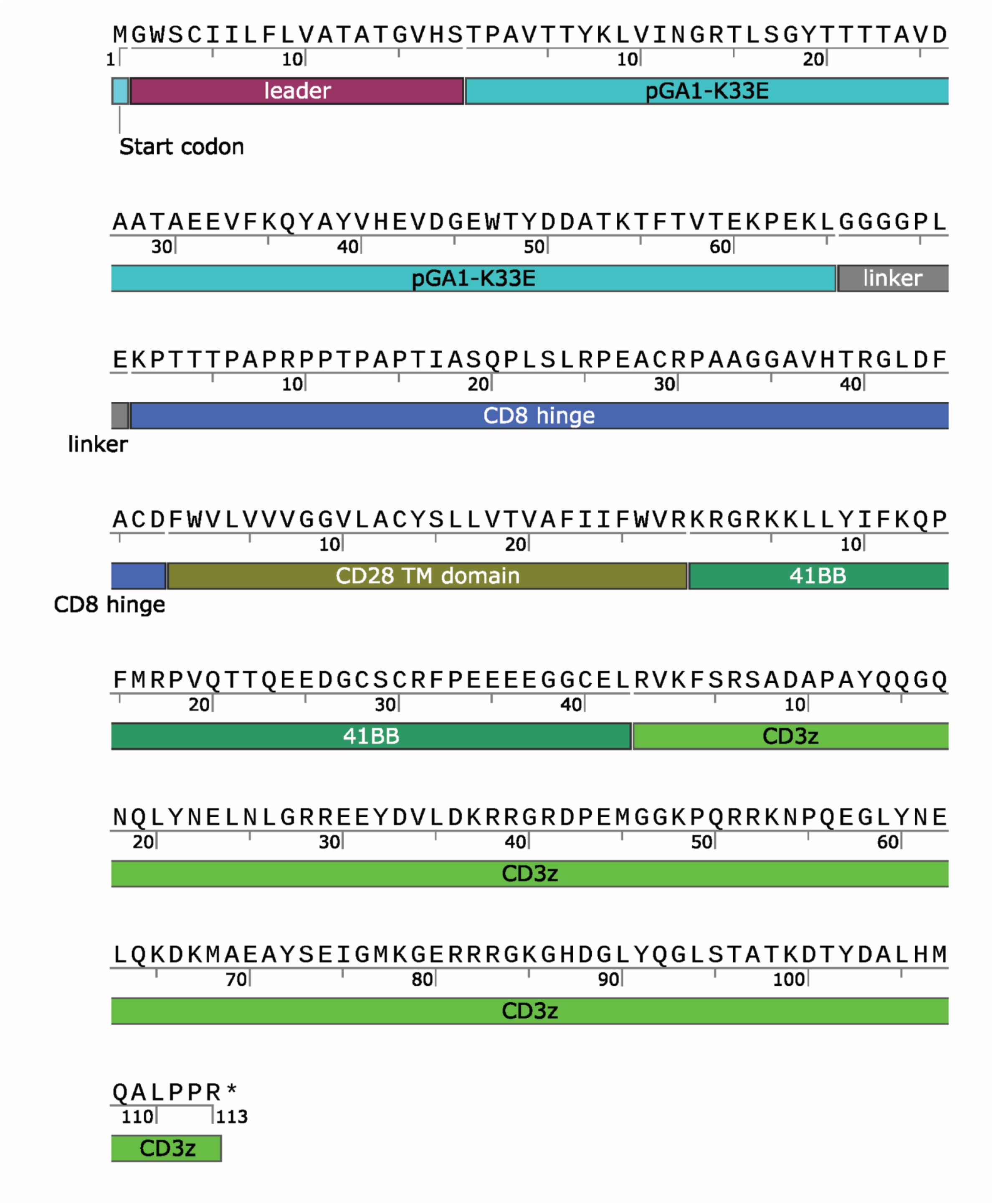
GA1CAR amino acid map sequence.

**Extended Data Fig 2.**
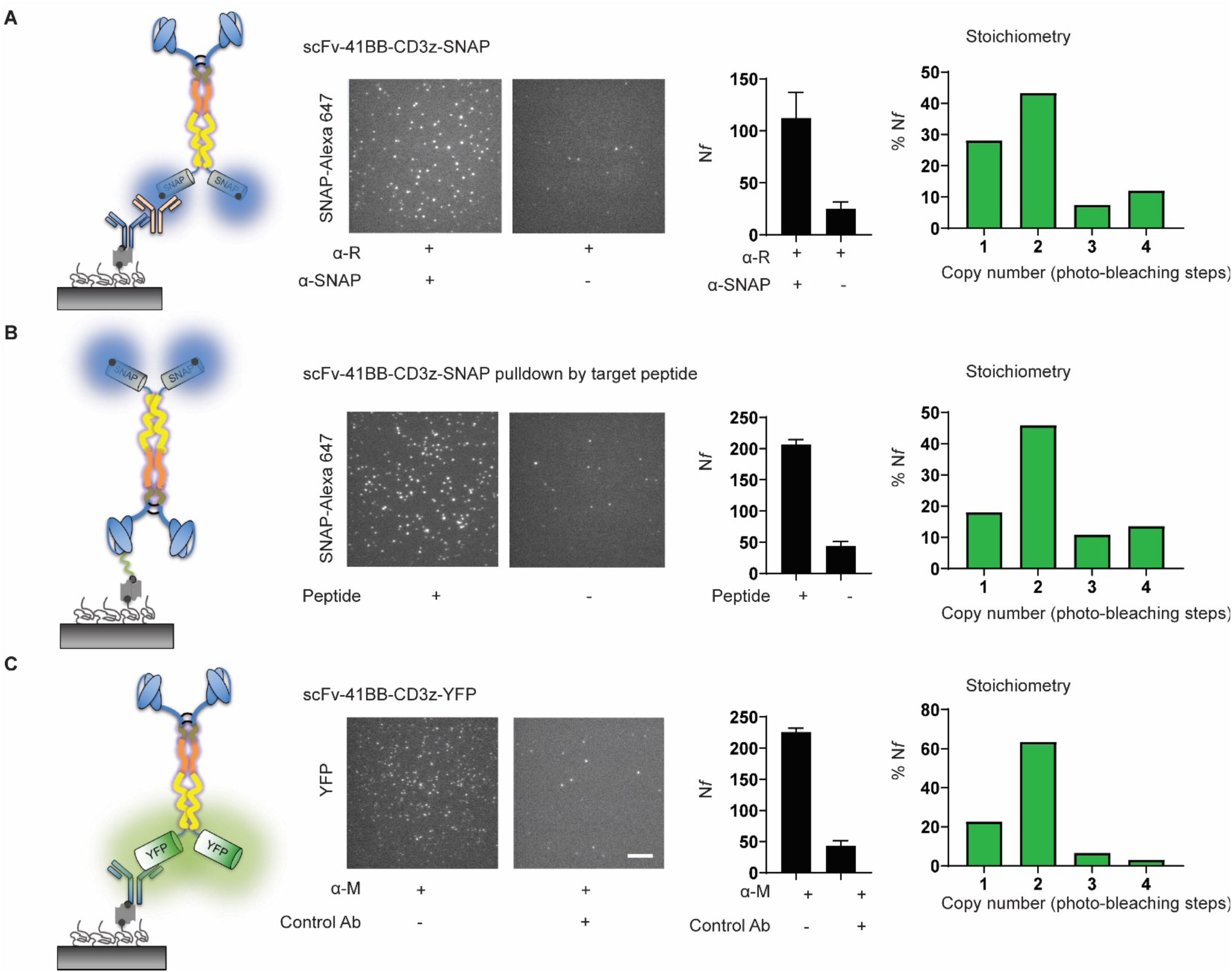
GA1CAR is dimeric. Schematic depiction of pulldown, representative fluorescent images, average number of molecules per imaging area (N*f*), and distribution of fluorescent photo-bleaching steps (stoichiometry) from A) CAR-T pulldown via anti SNAP rabbit antibody and anti-R antibody; B) CAR-T pulldown via biotinylated peptide that bind to scFv in CAR-T; C) CAR-T pulldown via anti-YFP mouse biotinylated antibody. scale bar 5 µm.

**Extended Data Fig 3.**
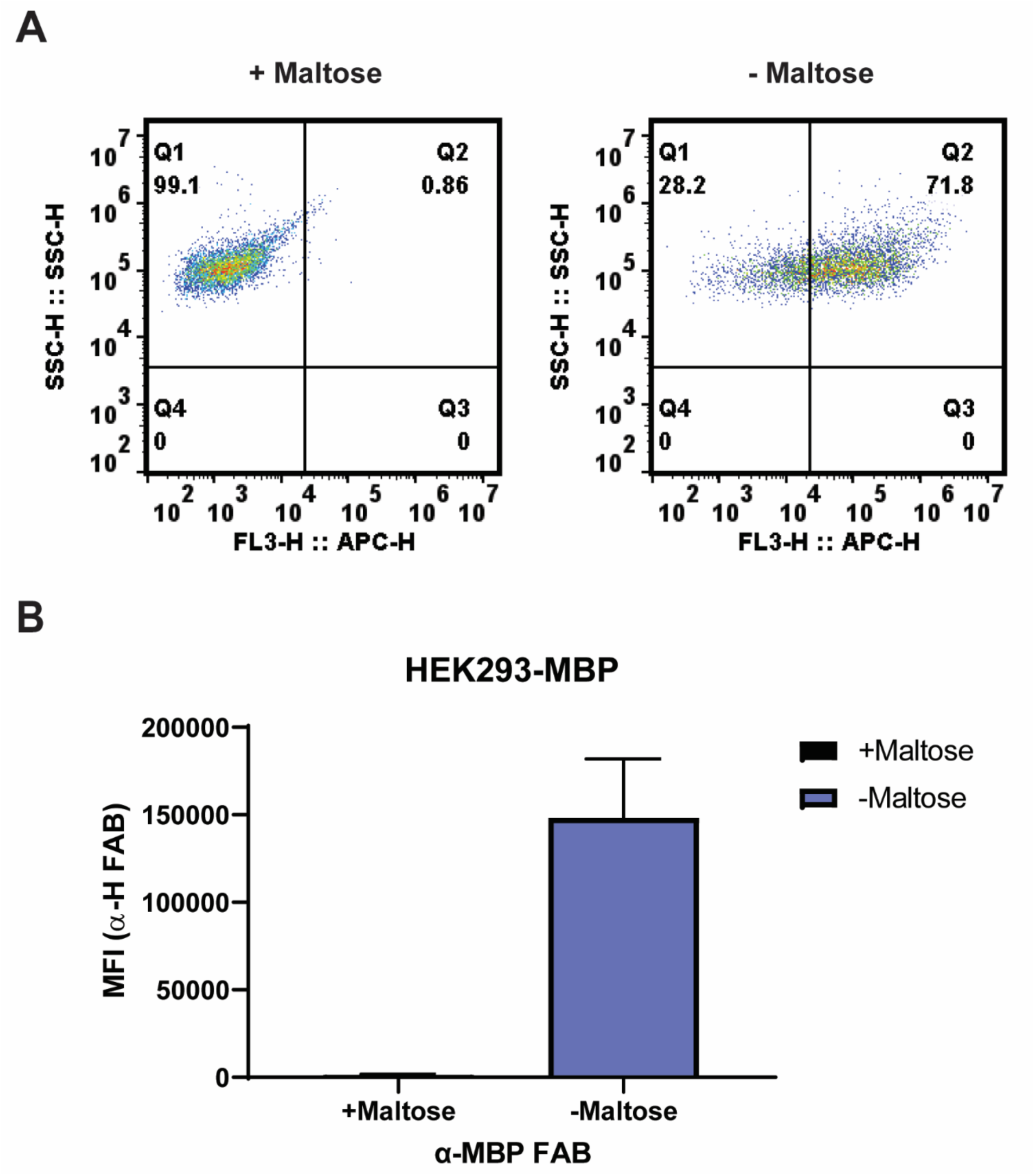
Expression of MBP on HEK-Flp cells. A) Representative flow plots showing expression of MBP on the surface of human embryonic kidney (HEK) 293 cells, detected using an anti-MBP Fab^LRT^ and a secondary anti-human Fab antibody conjugated to Alexa Fluor® 647. Binding of conformation-specific anti-MBP Fab^LRT^ to MBP decreases in the presence of maltose. B) Quantification of binding as determined by the mean fluorescent intensity (MFI). Anti-MBP Fab^LRT^ was used at 50 nM. The data are presented as the mean ± SD, n = 3.

**Extended Data Fig 4.**
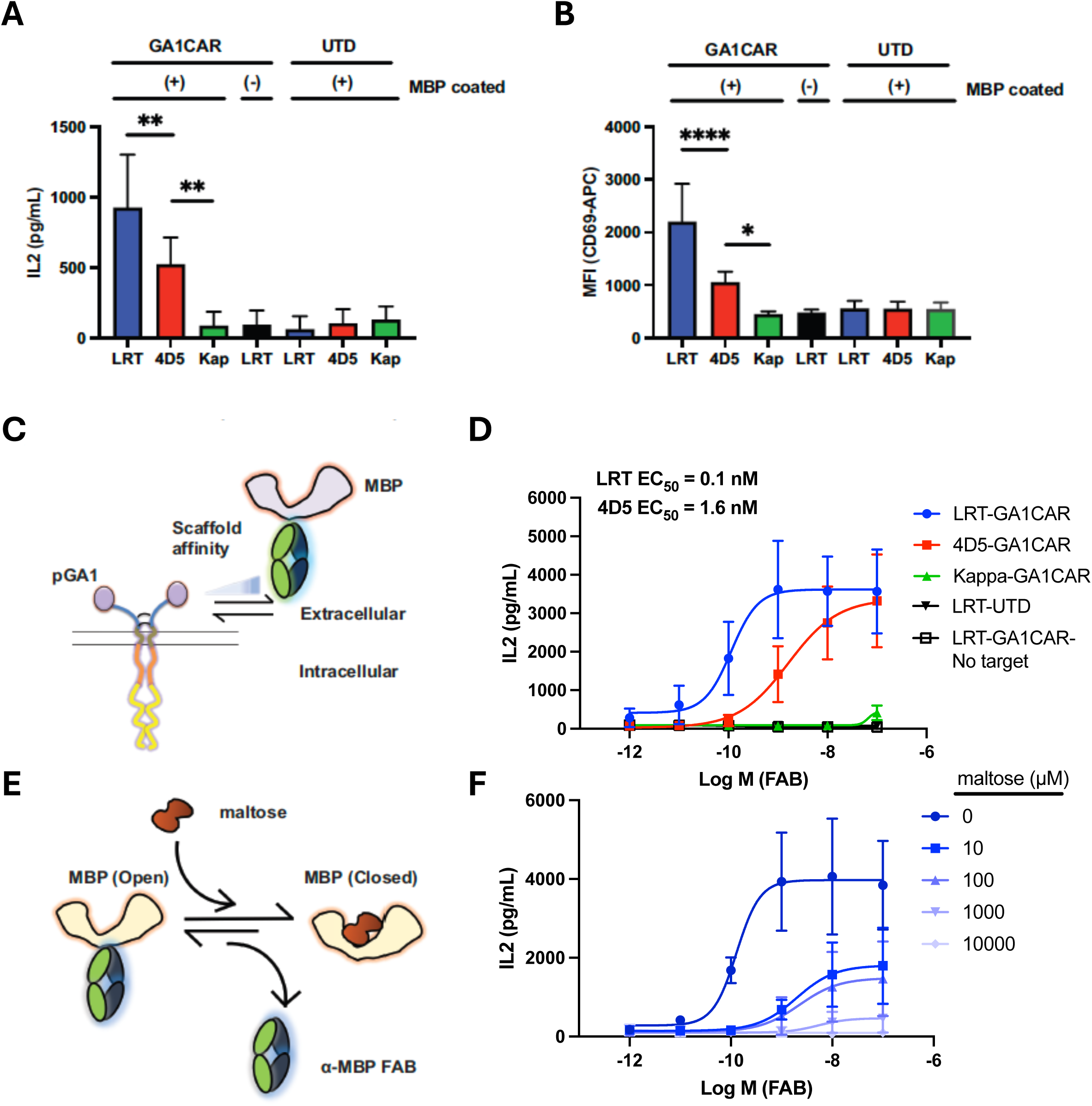
Functional characterization of GA1CAR expressed in Jurkat T cell lymphoma cells. A) IL2 production and B) CD69 expression by GA1CAR-Jurkat cells after being cultured in MBP coated plates and incubated with different anti-MBP Fab scaffolds for 16 hr. C) Cartoon showing Fab scaffolds with different affinities for GA1CAR. D) Quantification of IL2 release by GA1CAR-Jurkat cells after 16-hour co-culture with HEK- MBP cells at increasing concentrations of anti-MBP Fab scaffolds. E) Cartoon showing the conformation-specific anti-MBP Fab binding to MBP; the affinity decreases with increasing maltose concentration. F. Affinity dependent release of IL2 by GA1CAR-Jurkat cells in the presence of HEK-MBP cells and varying concentrations of maltose. Statistical significance was determined by Tukey’s multiple comparisons test after one-way ANOVA (*P < 0.05; **P < 0.01; ***P < 0.001; ****P < 0.0001), P < 0.05 considered significant. The data are presented as the mean ± SD, n = 3.

**Extended Data Fig 5.**
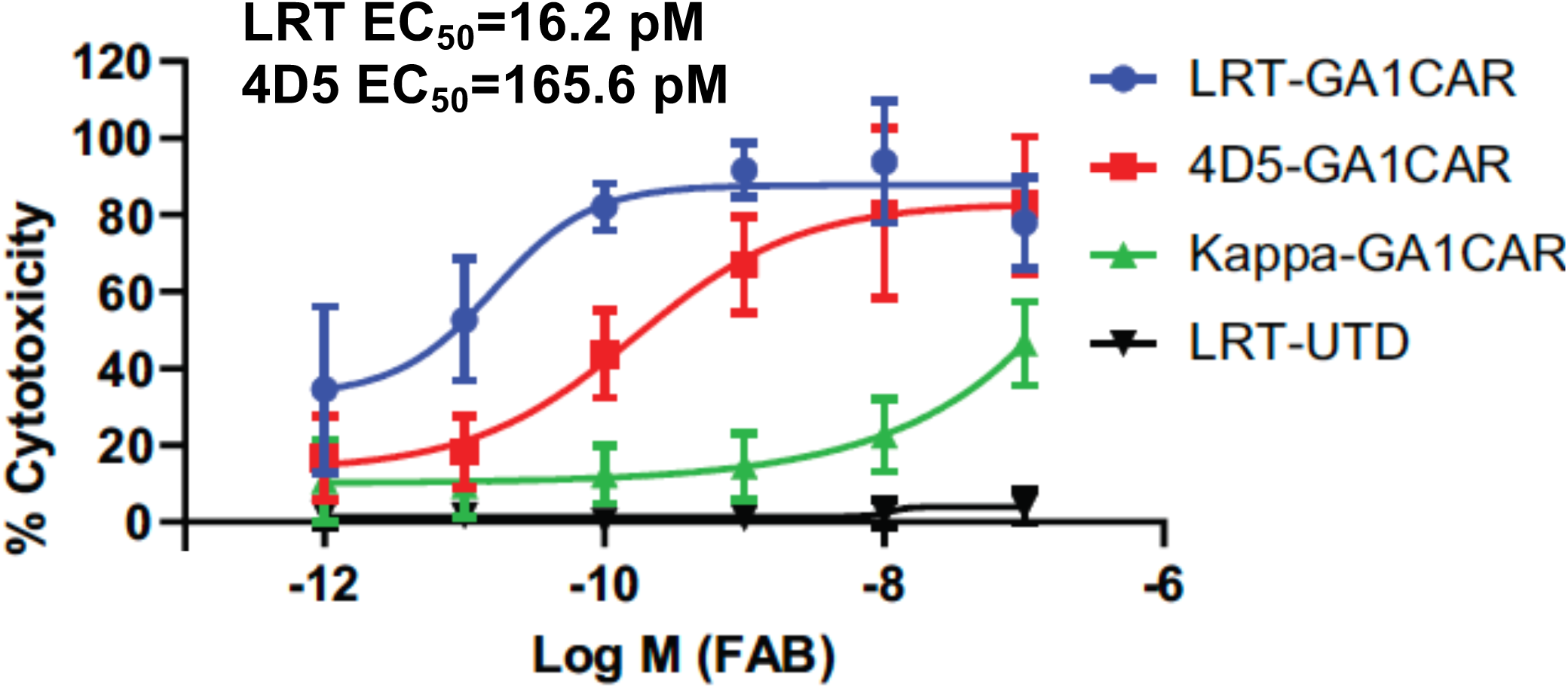
GA1CAR-T cell cytotoxicity when incubated with different concentration of anti-MBP Fab scaffolds and HEK-MBP cells. The data are presented as the mean ± SD, n = 3.

**Extended Data Fig 6.**
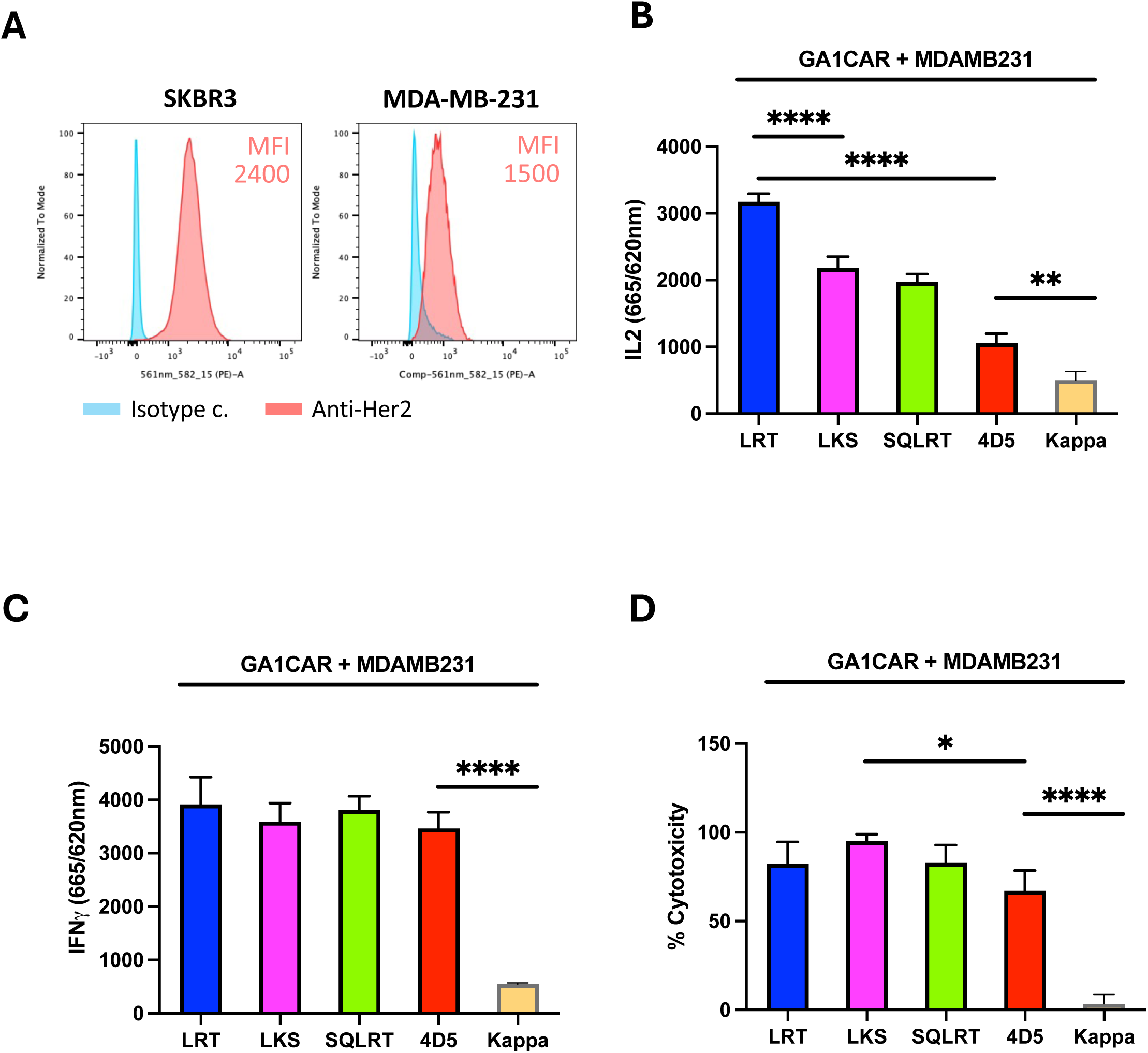
GA1CAR-T cell function using anti-Her2 Fab scaffolds with different affinities for GA1CAR and MDA-MB-231 cells as targets: A) Her2 expression is lower in MDA-MB-231 as compared to SKBR3 cells. B) IL2 release , C) IFNγ release, and D) Cellular cytotoxicity when incubating GA1CAR-T cells with MDA-MB-231 cells. Statistical significance was determined by Tukey’s multiple comparisons test after one-way ANOVA (*P < 0.05; **P < 0.01; ***P < 0.001; ****P < 0.0001), P < 0.05 considered significant. The data are presented as the mean ± SD, n = 3.

**Extended Data Fig 7.**
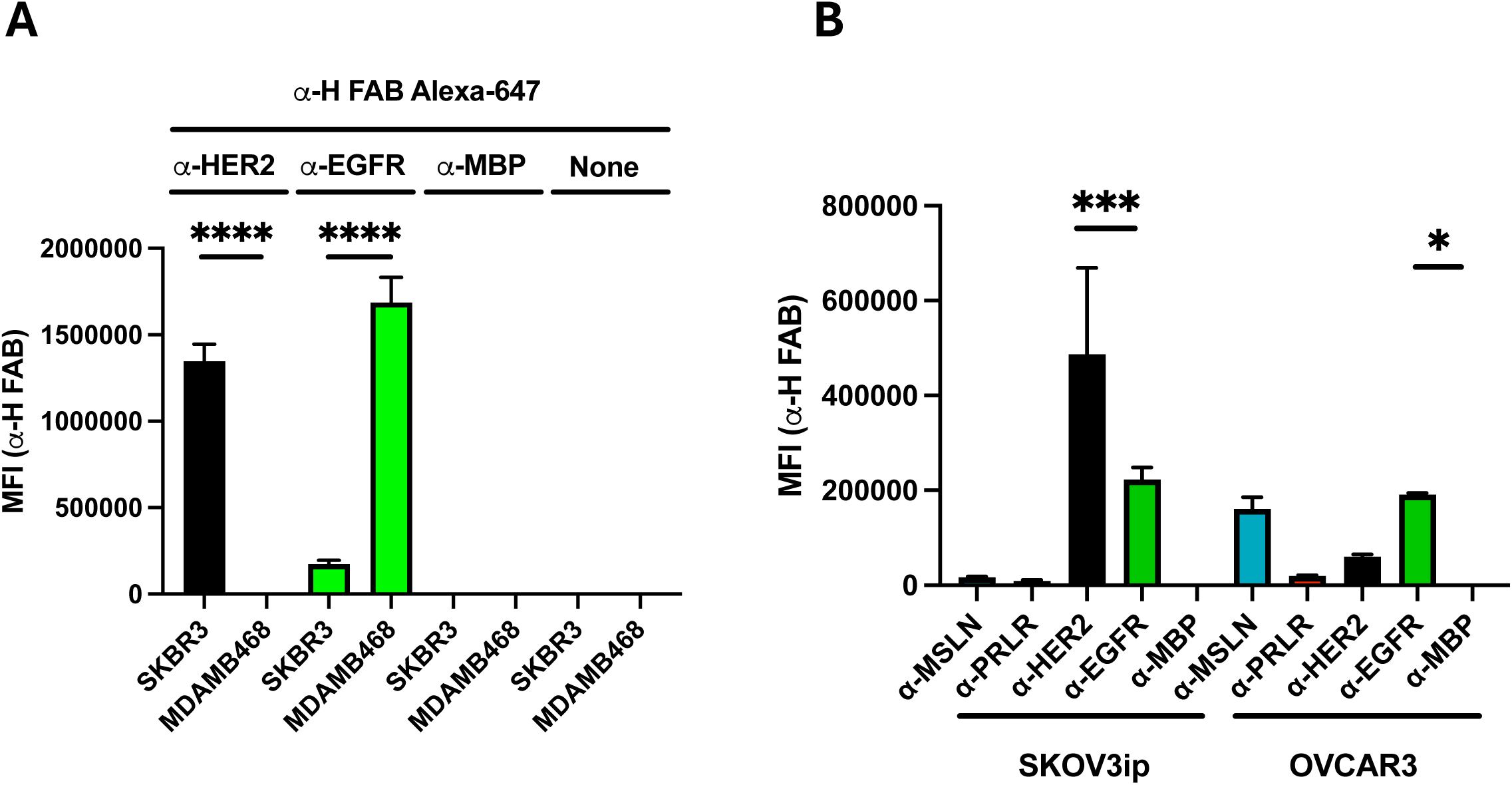
Quantification of cancer antigen expression on different target cells. A) Expression of HER2 and EGFR in SKBR3 and MDAMB468 breast cancer cell lines, determined by flow cytometry. B) Quantification of several ovarian cancer antigens on SKOV3ip and OVCAR3 cells by flow cytometry. Statistical significance was determined by Tukey’s multiple comparisons test after one-way ANOVA (*P < 0.05; **P < 0.01; ***P < 0.001; ****P < 0.0001), P < 0.05 considered significant; ns, no significant. The data are presented as the mean ± SD, n = 3.

**Extended Data Fig 8.**
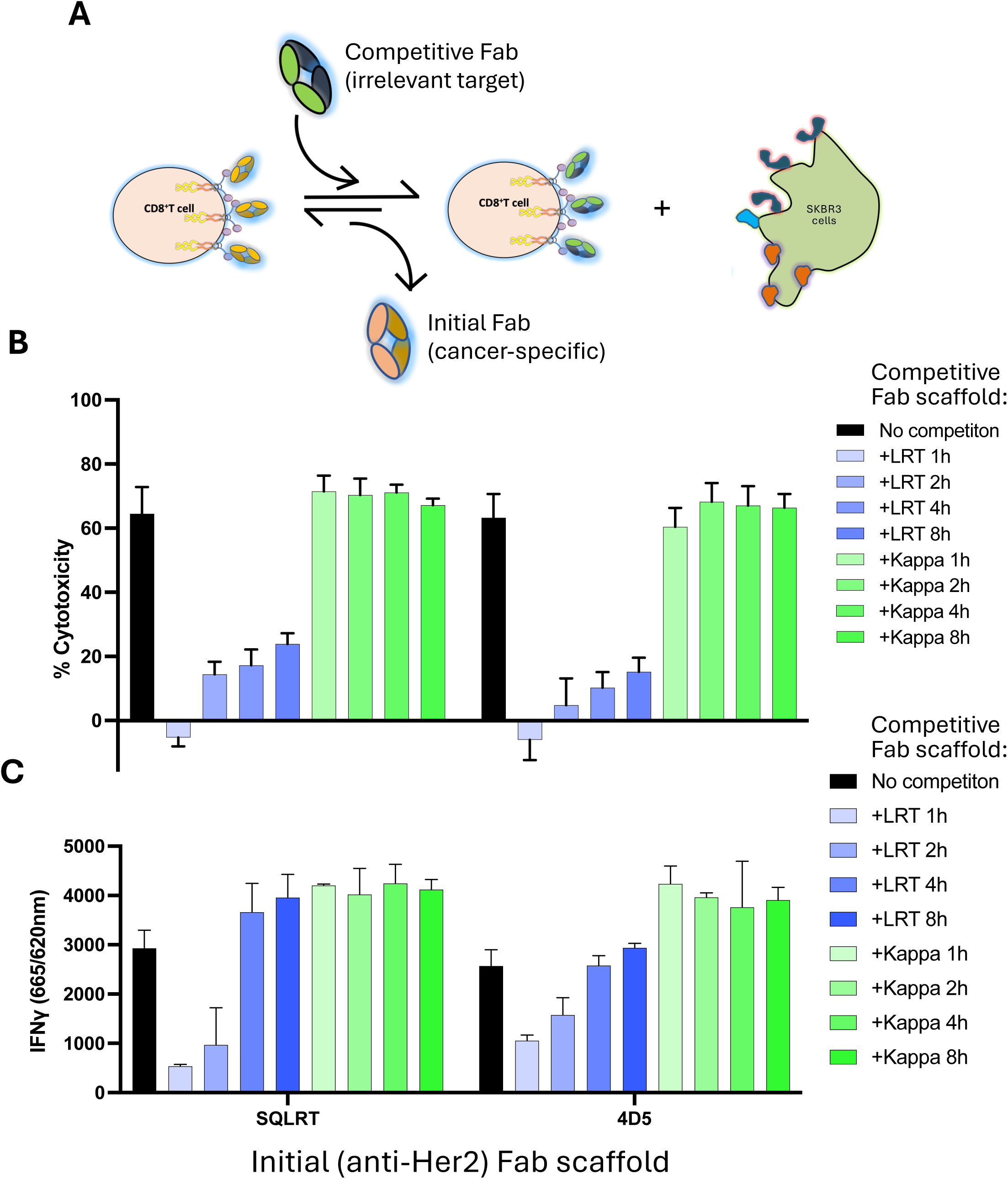
GA1CAR-T emergency stop switch. (A) Model of the GA1CAR-T emergency stop switch. A competitive isotype Fab scaffold with a higher affinity to GA1 can compete and deactivate the GA1CAR-T cells. Her2^+^ SKBR3 cancer cells were attached to the plate overnight, followed by addition of the GA1CAR-T cells and 100 nM of HER2 Fab in either SQLRT or 4D5 scaffolds the next day. Cell killing detected by LDH assay (B), and IFNγ release (C) were measured after 24h. To test the effect of the emergency stop switch, 1 μM of the competitive isotype Fab in the LRT scaffold was added after 1h, 2h, 4h or 8h later (isotype Fab recognizes SARS-CoV-2 RBD protein). The same isotype Fab in the Kappa scaffold was added as a negative control. We observed a reduction of cell killing upon CAR-T detachment from the HER2 Fabs in both SQLRT and 4D5 scaffolds (blue bars) compared to no emergency switch off (black bar) or control Kappa scaffold (green bar). The data are presented as the mean ± SD, n = 3.

**Extended Data Fig 9.**
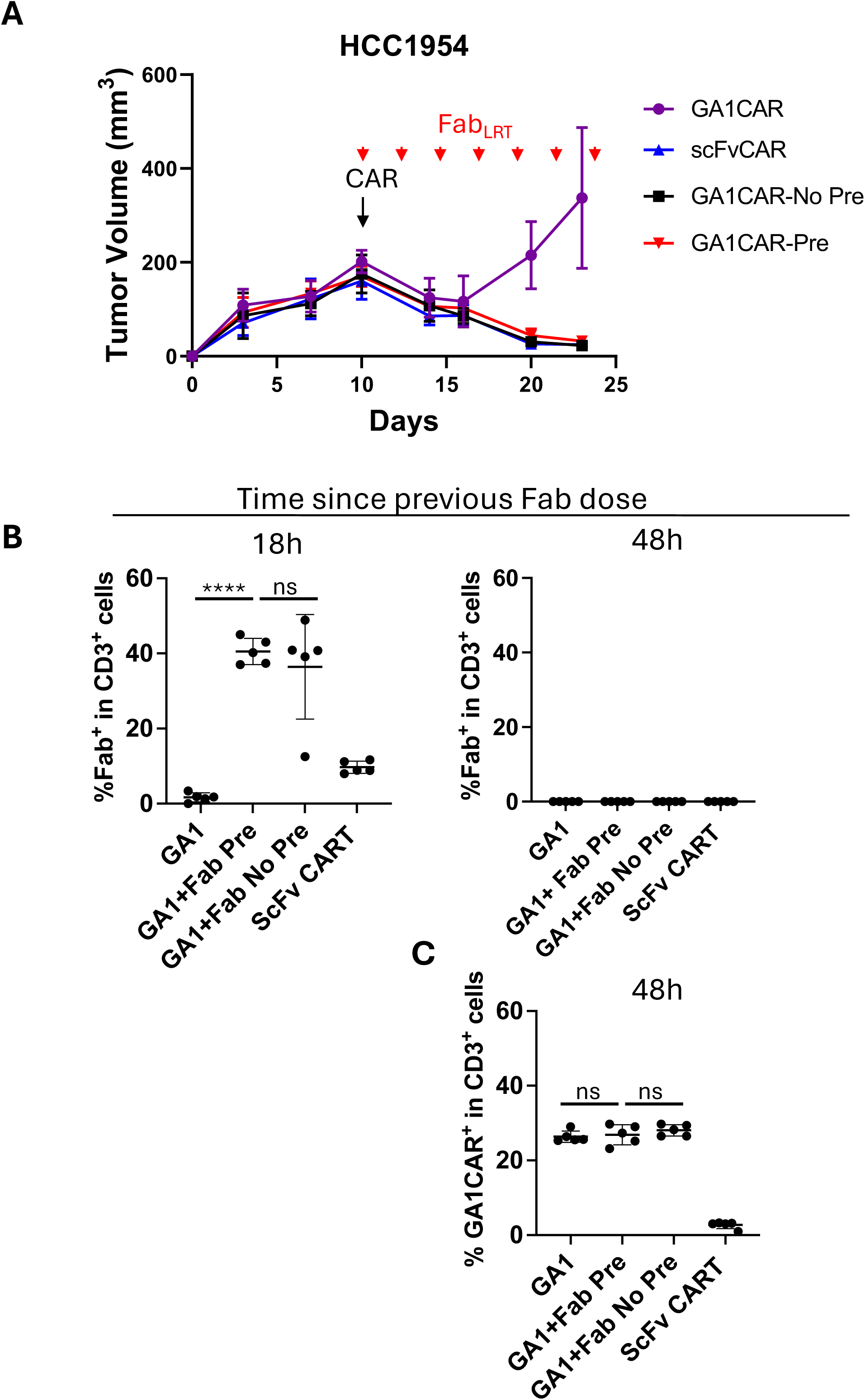
Pre-incubation with Fab^LRT^ is not required for effectiveness of GA1CAR-T cells in vivo and kinetics of Fab^LRT^ bound to circulating GA1CAR-T cells. HCC1954 tumor-bearing mice were injected i.v. at day 10 with GA1CAR-T cells that had been preincubated with HER2 Fab^LRT^ for 30 min before injection or not preincubated. The latter group of mice received Fab^LRT^ i.p at day 10. Both groups received 6 additional i.p. doses of Fab^LRT^ given every other day. Control groups included mice treated with GA1CAR-T cells but no Fab^LRT^, and mice treated with scFv CAR-T cells (N=5/group). B- C) Mice were bled 18 h after receiving the first dose of Fab^LRT^ and 48h after the third dose, right before treatment with a fourth dose. B. PBL were directly stained with an anti-human F(ab’)2 antibody to detect Fab^LRT^ bound to GA1 in vivo. C. PBL were preincubated with an irrelevant Fab^LRT^ and then stained with an anti-human Fab antibody to measure GA1 expression, 48 h after the previous Fab^LRT^ treatment. Statistical significance was determined by one-way ANOVA followed by Šídák’s multiple comparison tests (****P < 0.0001), ns: not significant. The data are presented as the mean ± SD, n = 5.

**Extended Data Fig 10.**
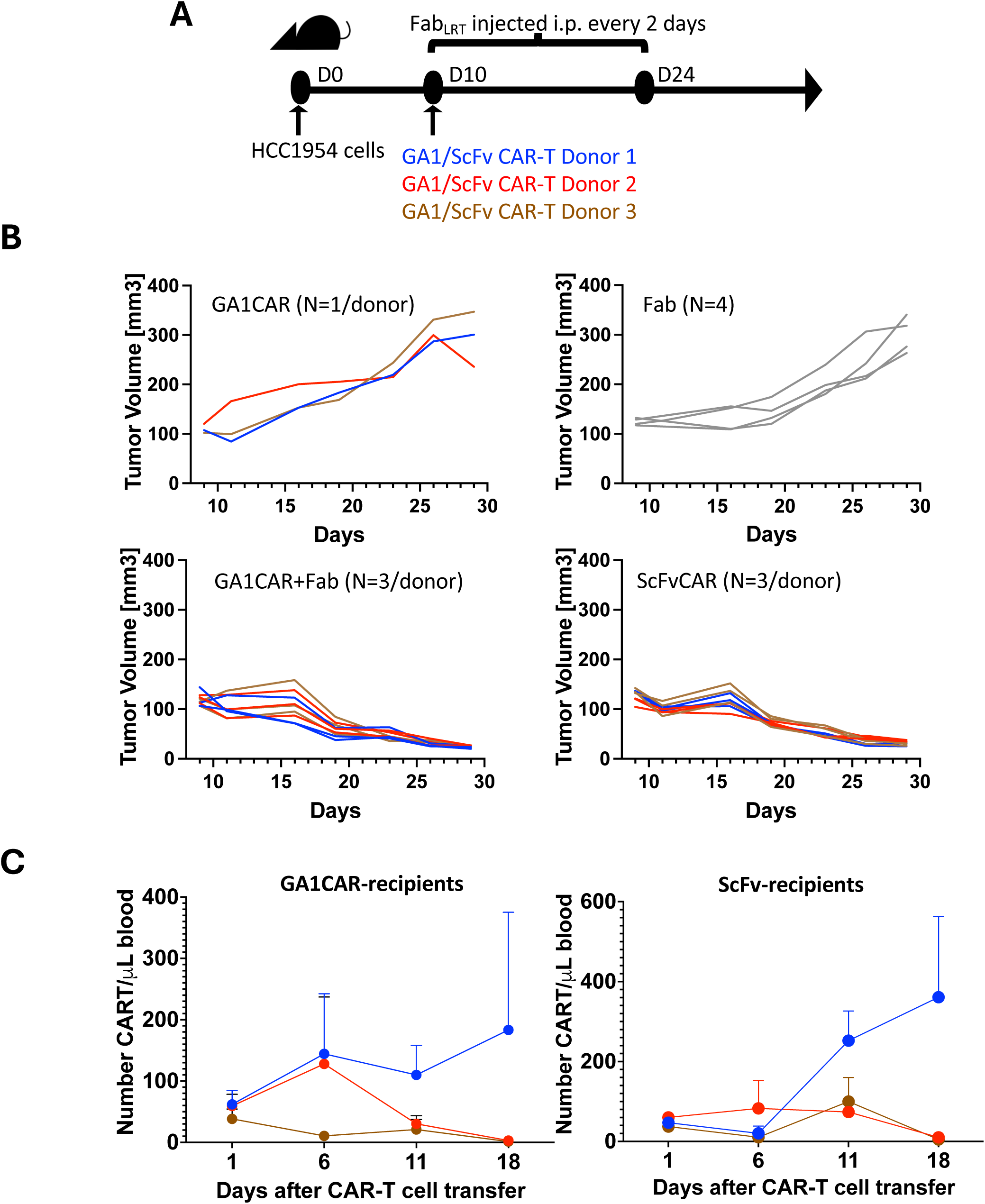
Kinetics of circulating CAR-T cells and of tumor elimination are similar for mice treated with GA1 and scFv CAR-T cells. A. Experimental scheme. HCC1954-bearing NSG mouse cohorts were treated with donor-matched GA1+Fab^LRT^ or scFv CAR-T cells, from three independent donors. Control groups received only GA1CAR-T cells or only Fab^LRT^. B. Tumor growth. Each line is an individual mouse, and colors (blue, red, brown) indicate different donors. C. Absolute numbers of circulating CAR-T cells were determined at different time points after CAR-T cell transfer, for each CAR-T cell type and donor. The data are presented as the mean ± SD, n = 3 recipients per donor and time point.

**Extended Data Fig 11.**
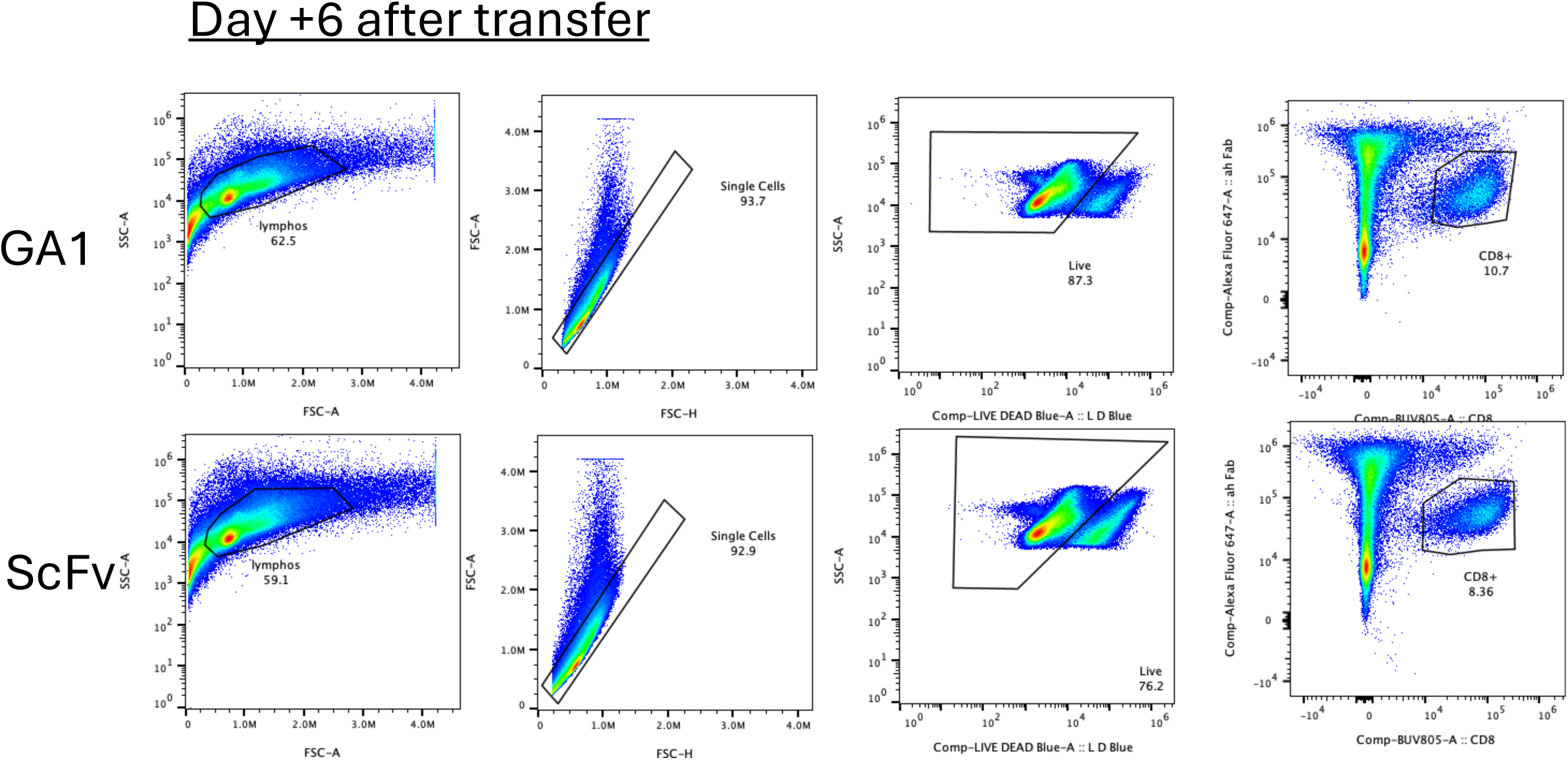
Gating strategy for quantification of tumor-infiltrating GA1 and scFv CAR-T cells by flow cytometry, and for sorting of populations for RNAseq of ex vivo isolated samples.

**Extended Data Fig 12.**
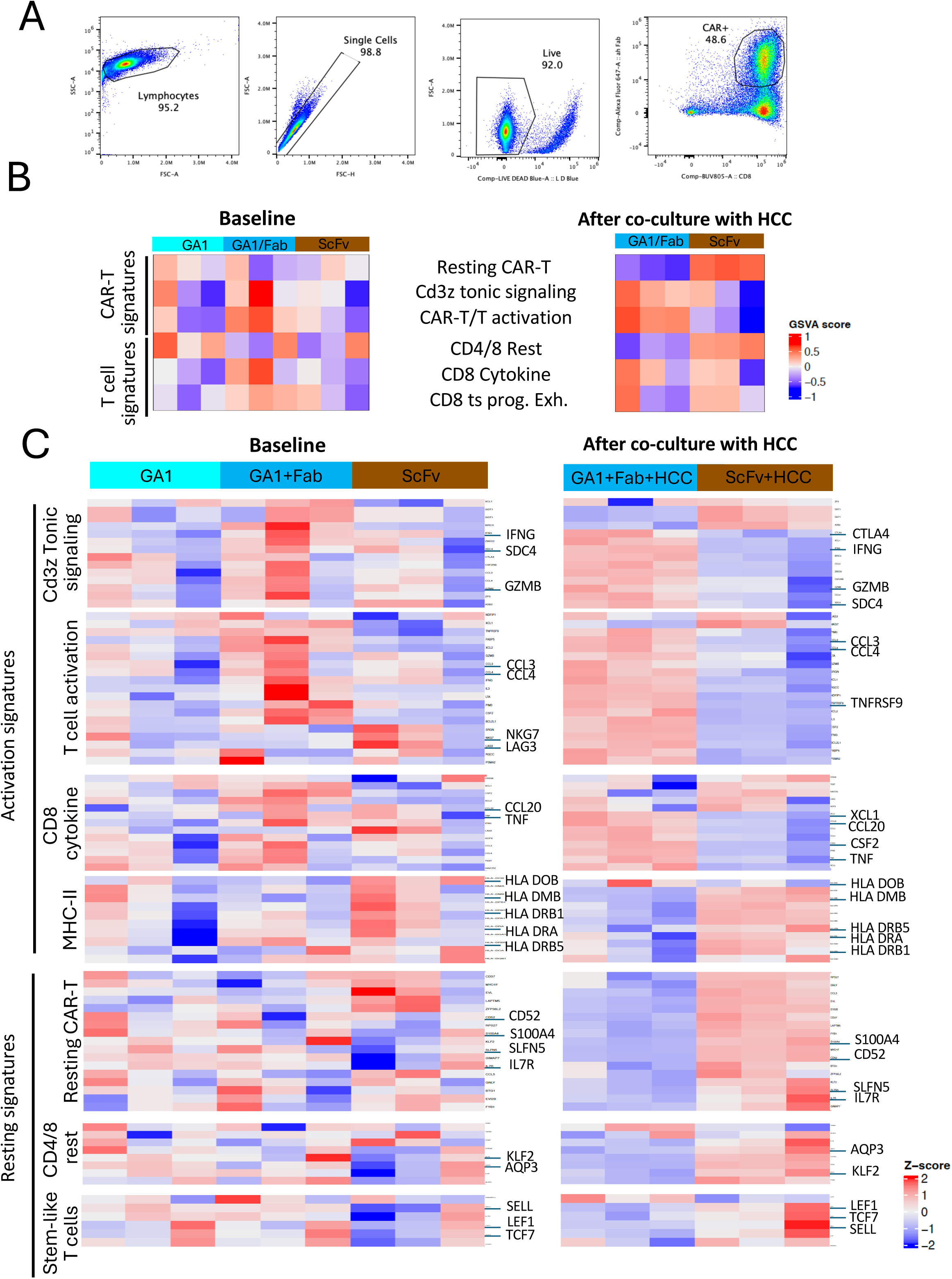
RNAseq data from in vitro activated GA1 and scFv CAR-T cells. A. Gating strategy for sorting of GA1 and scFv CAR-T cells cultured in vitro. A single representative GA1 sample is shown here, but GA1 and scFv gates from all donors can be seen in Extended Data Fig 13. B-C. Enrichment in gene signatures indicative of resting and activated CAR-T and T cell status was compared between scFv and GA1CAR-T cell samples from three independent donors. B) GSVA scores for each signature, C) Genes used for each gene signature, with representative genes highlighted.

**Extended Data Fig 13.**
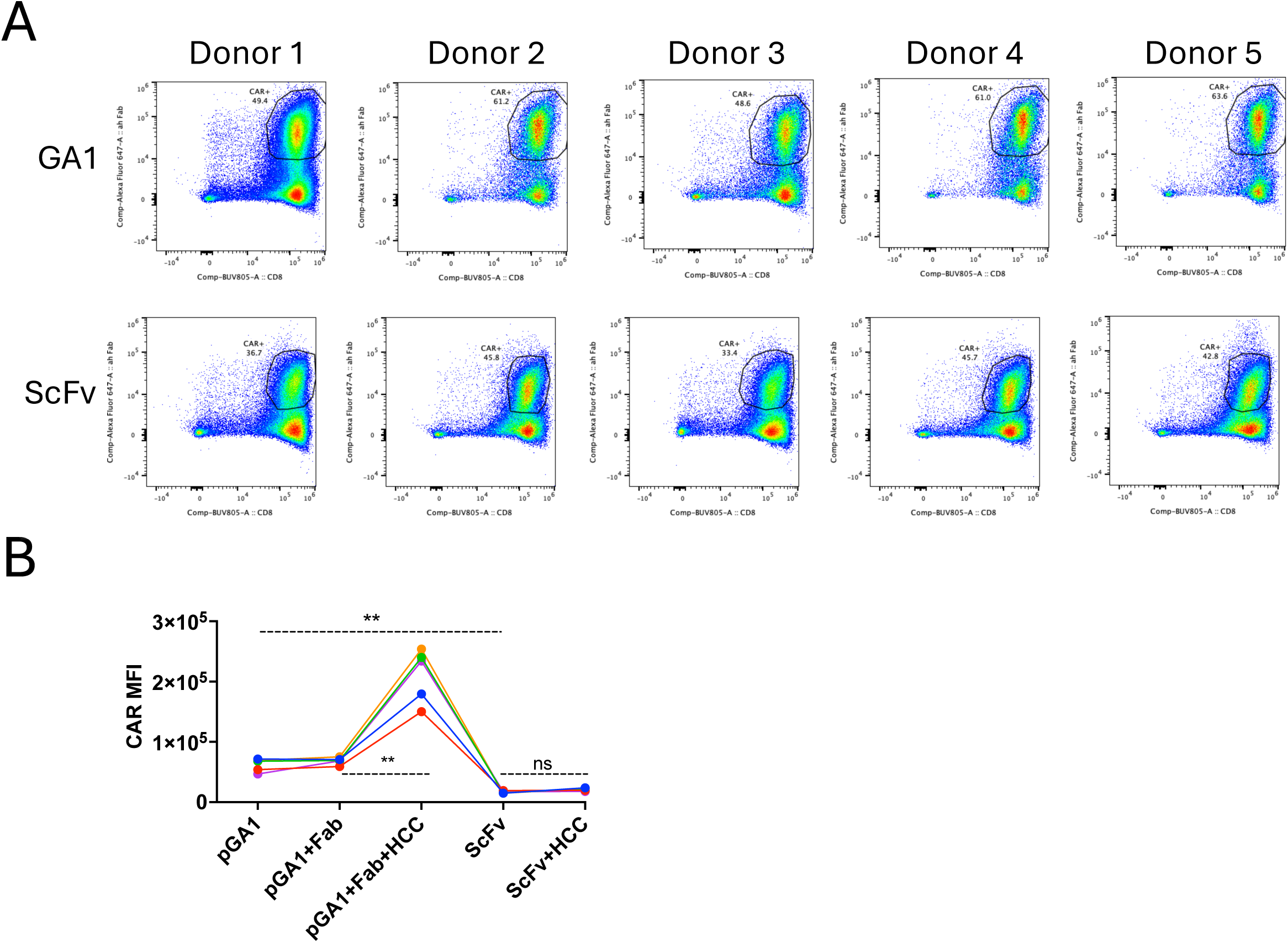
Variations of CAR expresion on CAR-T cell surface according to CAR type and donor (A) and experimental condition (B). Statistical significance on B. was analyzed using repeated measurements ANOVA followed by Šídák’s multiple comparison tests (**P < 0.01), ns:not significant. The data are presented as the mean ± SD, n = 5.

